# Curcumin–Gold Nanocomposites for Enhanced Cationic Anticancer Drug Delivery: Molecular Mechanisms of Loading and Membrane Interactions

**DOI:** 10.64898/2026.05.18.725887

**Authors:** Anuj Garg, Subodh Barik, Harikrishna Nair, Sreedev G. Nair, J. K. Kiran Kumar, Subbarao Kanchi

## Abstract

Curcumin-functionalized gold nanoclusters are promising platforms for catalysis and drug delivery, however, the molecular determinants governing their stability, morphology, and solvent response remain poorly understood. Here, microsecond all-atom molecular dynamics simulations were employed to investigate a 2 nm gold nanoparticle noncovalently coated with different curcumin forms, including the neutral enol and trans-keto tautomers, the deprotonated enolate, and their mixtures in water-ethanol and water-methanol solvents. Region-resolved analyses of the radius of gyration, density profiles, and surface coverage reveal that neutral enol and trans forms generate compact assemblies with near-complete surface coverage, whereas enolate-rich systems adopt more expanded conformations with greater solvent exposure. Mixed systems preserve these intrinsic packing characteristics while improving overall surface coverage. Solvent substitution from ethanol to methanol reduces π-π stacking, strengthens Au-curcumin interactions, and increases surface coverage, yielding more compact nanostructures. Free energy and potential of mean force calculations further indicate that deprotonated curcumin most effectively screens Au-Au interactions and promotes nanoparticle dispersion, whereas the neutral tautomers provide moderate stabilization. Curcumin functionalization also enhances the loading of the anticancer drugs doxorubicin (DOX) and niraparib (NIR) onto Au nanoparticles. Membrane interaction with Au-DOX/NIR-Curcumin simulations show that Enolate(An)-containing systems form more extended structures and interact more weakly with both negatively charged and neutral DMPC membranes. In contrast, neutral curcumin complexes form compact, positively charged assemblies with stronger interactions with the negatively charged model membrane. Furthermore, these interactions are predominantly driven by electrostatic attraction and are substantially weaker with the neutral DMPC membrane. Overall, these findings demonstrate how curcumin tautomeric state and solvent environment cooperatively govern interfacial organization and colloidal stability, drug-loading, and membrane interactions. This provides molecular-level design principles for curcumin-based gold nanocarriers for catalysis, sensing, and drug delivery applications.

## Introduction

Functionalization of ultrasmall gold nanoparticles (1–3 nm in diameter) with organic ligands affords precise control over size, surface chemistry, and colloidal stability, thereby enhancing catalytic performance and imparting distinct, molecule-like optical and electronic characteristics that underpin their widespread use in materials synthesis and biomedical applications (*1–4*). Their atomically precise compositions, commonly denoted as [*M_n_L_m_*]*^q^*, enable rigorous structure– property correlations that are crucial for rational design in catalysis, sensing, and biomedicine (*5–8*). The synthesis of gold nanostructures for applications such as vitamin sensing and the catalytic reduction of nitro-compounds can be strongly influenced by growth temperature and pH conditions, respectively (*9,10*). At the heart of these behaviours lies the metal–ligand interface, where ligand identity and binding mode govern nanoparticle stability, reactivity, solubility, and biological interactions (*11–15*). Ligand attachment to gold surfaces can be achieved through covalent strategies employing linker molecules such as thiols or PEG, or via noncovalent adsorption approaches that preserve the native ligand chemistry. Although covalent conjugation often provides superior thermodynamic stability, it can also increase toxicity and perturb the desirable physicochemical properties of the ligand shell. In contrast, noncovalent functionalization typically enables simpler synthesis, eliminates the need for additional stabilizing agents, and reduces cytotoxicity. Recent reports demonstrate that polymers and protein scaffolds can simultaneously serve as reducing agents and stabilizing ligands for gold nanoparticles, yielding highly luminescent, biocompatible constructs with exceptional optical properties (*16,17*). This dual-function strategy motivates broader exploration of bioorganic ligands beyond conventional thiols, including naturally derived molecules such as curcumin that can act as both reducing and templating agents during nanoparticle formation. Bioorganic ligands bearing aromatic moieties, hydrogen-bonding functionalities, and extended *π*-electron systems have consequently emerged as attractive alternatives to thiols and phosphines, offering tunable interfacial chemistry, improved biocompatibility, and versatility for advanced applications in catalysis and nanomedicine (*18–21*).

Curcumin, a polyphenolic β-diketone component of the common kitchen spice turmeric (*Curcuma longa*), features a conjugated π-system, dual phenolic –OH groups, and a β-diketone unit that readily chelates metal ions (*22–24*). These structural features drive its versatile reactivity and enable robust noncovalent binding via π–π stacking, hydrogen bonding, and hydrophobic interactions at nanoparticle interfaces (*25–27*). In addition to this chemical adaptability, curcumin’s extensively validated antioxidant, anti-inflammatory, antimicrobial, and anticancer properties (*28–31*) have fueled investigations into its metal complexes for therapeutics and sensors. Notably, its strong affinity for gold ions and ability to reduce Au precursors under ambient conditions make it an excellent agent for green, surfactant-free synthesis of gold nanostructures (*32–35*).

Curcumin-functionalized gold nanoparticles (Au–Cur NPs) constitute a distinctive subset of bioorganic ligand-modified gold nanostructures that leverage curcumin as both a green reducing agent and a stabilizing shell, thereby eliminating the need for additional toxic reductants and surfactants while yielding sustainable, biocompatible nanocarriers (*36–38*). In such “green” synthetic protocols, curcumin simultaneously mediates Au precursor reduction and nanoparticle passivation, providing size-controlled particles typically in the 18-40 nm range that support efficient cellular uptake and often incorporate pH-responsive release behavior tailored to the acidic tumor microenvironment (*39,40*). These site-selective release characteristics enhance therapeutic efficacy and limit off-target damage, essential for targeted cancer therapy (*41,42*). Conjugation of curcumin onto gold nanoparticle surfaces markedly improves its intrinsic aqueous solubility and systemic bioavailability. This enhancement enables higher effective intracellular concentrations and potentiated antiproliferative and pro-apoptotic responses in cancer cell lines, including MCF-7 (breast), HCT-116 (colon), C6 (glioma), and Huh7 (liver) (*36,39,43*). Furthermore, co-delivery strategies that co-load curcumin with additional bioactive agents such as sulforaphane have been shown to enhance anticancer efficacy by concurrently targeting multiple cellular pathways (*44,45*).

Beyond their chemotherapeutic performance, Au-Curcumin NPs exhibit pronounced theranostic potential by uniting drug delivery with optical and fluorescence-based imaging modalities derived from the plasmonic and emissive properties of the gold core (*46,47*). This multifunctionality enables real-time monitoring of drug distribution and treatment response, and can be further refined through the incorporation of targeting ligands, such as folate or cyclodextrin, to enhance tumor selectivity and minimize systemic side effects (*48–50*). In some formulations, Au– Cur nanostructures are capable of traversing the blood-brain barrier and expressing strong antioxidant and anti-inflammatory activities, thereby extending their relevance to neurodegenerative disorders, fibrosis, wound healing, and tissue-regeneration scenarios (*32,42,43,51,52*). These systems also act as enzyme inhibitors that can suppress deleterious processes such as amyloid fibril formation, underscoring their broader potential in complex pathologies beyond cancer (*43,53*). Collectively, the improved solubility, bioavailability, targeting capability, and pH-responsive release of curcumin achieved through Au–Curcumin architectures highlight them as versatile platforms whose experimentally observed performance motivates a deeper molecular-level understanding of their interfacial organization, stability, and solvent-dependent behavior – as addressed in the present investigation.

Doxorubicin is a commonly employed anticancer drug that inhibits tumor progression by inserting into DNA and blocking topoisomerase II activity, thereby interfering with replication and transcription processes (*54,55*). Loading doxorubicin onto gold nanoparticle carriers enables more precise and controlled delivery, potentially reducing side effects such as cardiotoxicity and bone marrow suppression (*56*). Previous studies have demonstrated that such nano-carrier-based systems can enhance the efficiency of targeted doxorubicin delivery up to higher doses while minimizing damage to surrounding healthy tissues (*56,57*). Furthermore, co-delivery of doxorubicin with curcumin within nanocarriers such as polymeric micelles provides additional therapeutic benefits, particularly in mitigating cardiotoxicity. Curcumin enhances the sensitivity of cancer cells to doxorubicin by modulating drug resistance pathways, including PI3K/Akt and NF-κB signaling, and by inhibiting P-glycoprotein activity (*54*). It also offers cardio-protective effects by reducing oxidative stress and limiting the accumulation of doxorubicinol, a metabolite associated with cardiac damage. Given its poor bioavailability, curcumin benefits from encapsulation along with doxorubicin and its derivatives, which improves its stability and systemic availability (*58*). This combined delivery system exhibits prolonged circulation time and enhanced accumulation at tumor site, thereby improving overall therapeutic efficacy.

Experimental studies have provided valuable insights into curcumin-functionalized gold nanoparticles. However, density functional theory (DFT) calculations elucidate the molecular level mechanisms governing their stability and functionality. In gelatin-coated AuNPs, charge transfer from gold core to gelatin matrix stabilizes the complex, explaining its high encapsulation efficiency and pH-responsive drug release behavior (*59*). Likewise, in curcumin-capped gold nanoparticles, disruption of intramolecular hydrogen bonds increases the accessibility of phenolic groups, thereby enhancing antioxidant potential (*32*). While DFT provides detailed static insights into electronic structure and energetics, molecular dynamics (MD) simulations capture dynamic solvent effects, thermal fluctuations, and ligand reorganization. Together, these methods provide a more comprehensive understanding of nanoparticle stability, curcumin orientation, and binding modes under physiologically relevant conditions. The existing literature does not provide a comprehensive understanding of the kinetic and thermodynamic processes involved in curcumin-mediated reduction of Au(III) species, particularly with respect to the roles of enolic tautomeric forms and solvent-dependent redox behaviour. This limited mechanistic insight also extends to the stabilization of curcumin on gold nanoclusters, where key factors including ligand-cluster interfacial dynamics, conformational flexibility, and solvent-mediated effects, remain insufficiently characterized. Clarifying these interconnected mechanisms is therefore essential for the rational design of synthesis strategies and for improving the reproducibility and functional reliability of curcumin-functionalized nanocarriers. To address these gaps, the structure, dynamics, and stability of Au-curcumin systems as a function of different solvent environments are investigated using all-atom molecular dynamics simulations. This study is further extended to evaluate the role of Au-curcumin composites in enhancing the loading of doxorubicin (DOX) and niraparib (NIR) onto Au nanoparticles. Accordingly, DOX/NIR complexation with Au-curcumin systems is examined under both ethanol and ethanol-free conditions. In addition, interactions of Au+DOX/NIR+curcumin nanocomposites with cancer cell membranes are critical for effective drug delivery and reducing side effects. Therefore, MD simulations were performed to investigate the adsorption behaviour of these complexes on a negatively charged model cancer-cell membrane, with a neutral DMPC membrane serving as a control, with particular emphasis on the role of the curcumin form. Overall, this integrated study provides molecular-level insights into how the curcumin state governs the colloidal stability of Au-curcumin nanocomposites and their effectiveness as carriers for cationic anticancer drugs, including doxorubicin (DOX) and niraparib (NIR).

## Methodology

The noncovalent conjugation of gold nanoparticle with various forms of curcumin (enol, keto (Trans), and enolate (An)) was investigated, with system details provided in **Table 1**. A charge neutral spherical gold nanoparticle (236 atoms, 2.0 nm diameter) and DMPC and cancer membranes were constructed using CHARMM-GUI (*60*). Lennard-Jones (L-J) parameters for gold atoms were adopted from a previous study (*61*). The cancer cell membrane model comprised 512 lipids (256 per leaflet) and consisted of a physiologically relevant mixture of 1-palmitoyl-2-oleoyl-sn-glycero-3-phospho-L-serine (POPS), cholesterol (CHL), palmitoyl sphingomyelin (PSM), 1-palmitoyl-2-oleoyl-sn-glycero-3-phosphocholine (POPC), and 1-palmitoyl-2-oleoyl-sn-glycero-3-phosphoethanolamine (POPE). Detailed membrane composition is provided in **Table 9**, and the lipid to cholesterol ratio was maintained in accordance with previously reported studies (*62,63*). The Lipid21 FF was implemented for the lipids and cholesterol molecules. Curcumin, doxorubicin, niraparib, and ethanol structures were optimized using the hybrid functional comprising of the Becke’s exchange functional, Lee-Yang-Parr correlation functional and the Hartree-Fock exchange term (B3LYP) with a 6-311++G(d,p) basis set in Gaussian16, yielding partial atomic charges (**Table S1, Table S2, and Table S3**) from electrostatic potential (ESP) fitting (*64*). GAFF force field parameters for curcumin, doxorubicin, niraparib, and ethanol were obtained using Antechamber from the AmberTools package (*65*). Fifty curcumin molecules were randomly positioned around the gold nanoparticle avoiding steric clashes within a cubic simulation box of 9.85 × 9.85 × 9.85 nm³. The Au-curcumin systems were solvated in TIP3P water, with ethanol added to achieve 1:3 volumetric ratios. Systems were neutralized using H_3_O^+^ counter ions, followed by HCl addition to reach 0.1 M concentration, mimicking experimental conditions (*66–68*). The H_3_O^+^ counter ions were represented using the hydronium ion hydration free-energy (HFE) force field parameters (*69*). The force field parameters for Na^+^, Cl^−^ ions, and the Au representation are listed in **Table 10**. Similarly, Au and Au-curcumin nanoparticles loaded with doxorubicin/niraparib were prepared in the presence of ethanol under the conditions comparable to those described above. To investigate the effect of the solvent on the system, the simulations were further extended for an additional 50 ns in the absence of ethanol. The resulting final structures of the Au+DOX and Au+DOX+curcumin nanocomposites were subsequently positioned near the cancer membrane surface to elucidate their binding behaviour in the presence and absence of curcumin. These solvated systems then underwent microsecond all-atom molecular dynamics (MD) simulations in GROMACS 2024.4 using the protocol outlined below (*70*).

**Table 1:** Abbreviations list and Number of atoms information for all the systems prepared for the MD simulations (ETH – ethanol; MTH – methanol; SOL – water; mols – number of molecules).

| Abbreviations |  |
| --- | --- |
| An | Curcumin in the enolate form |
| Enol | Curcumin in the enol form |
| Trans | Curcumin in the trans keto form |
| DOX | Doxorubicin drug molecule |
| NIR | Niraparib drug molecule |
| An-Enol | Mixture of curcumin in the enolate and enol forms |
| Trans-An | Mixture of curcumin in the trans keto and enolate forms |
| Trans-Enol | Mixture of curcumin in the trans keto and enol forms |
| Au+An | Enolate form of curcumin surrounding gold nanoparticle |
| Au+Enol | Enol form of curcumin surrounding gold nanoparticle |
| Au+Trans | Trans keto form of curcumin surrounding gold nanoparticle |
| Au+An-Enol | Enolate and Enol forms of curcumin surrounding gold nanoparticle |
| Au+Trans-An | Trans keto and Enolate forms of curcumin surrounding gold nanoparticle |
| Au+Trans-Enol | Trans keto and Enol forms of curcumin surrounding gold nanoparticle |

| MD Simulation Information |  |  |  |  |  |  |  |  |  |
| --- | --- | --- | --- | --- | --- | --- | --- | --- | --- |
| S. No. | Au+curcumin Systems | Au | CUR | ETH (mols) | Charge | SOL (mols) | H<sub>3</sub>O<sup>+</sup> | Cl<sup>-</sup> | Total |
| 1 | Au+An | 236 | 2300 | 3000 | -50e | 28728 | 122 | 72 | 115914 |
| 2 | Au+Enol | 236 | 2350 | 3000 | 0 | 28828 | 72 | 72 | 116214 |
| 3 | Au+Trans | 236 | 2350 | 3000 | 0 | 28835 | 72 | 72 | 116235 |
| 4 | Au+An-Enol | 236 | 2325 | 3000 | -25e | 28847 | 97 | 72 | 116271 |
| 5 | Au+Trans-An | 236 | 2325 | 3000 | -25e | 28853 | 97 | 72 | 116289 |
| 6 | Au+Trans-Enol | 236 | 2350 | 3000 | 0 | 28852 | 72 | 72 | 116286 |
| 7 | An | --- | 2300 | 3000 | -50e | 28728 | 122 | 72 | 115678 |
| 8 | Enol | --- | 2350 | 3000 | 0 | 28828 | 72 | 72 | 115978 |
| 9 | Trans | --- | 2350 | 3000 | 0 | 28835 | 72 | 72 | 115999 |
| 10 | An-Enol | --- | 2325 | 3000 | -25e | 28847 | 97 | 72 | 116035 |
| 11 | Trans-An | --- | 2325 | 3000 | -25e | 28853 | 97 | 72 | 116053 |
| 12 | Trans-Enol | --- | 2350 | 3000 | 0 | 28852 | 72 | 72 | 116050 |
|  | Au+Drug+curcumin Systems | Au | CUR+ Drug (mols) | ETH (mols) | Charge | SOL (mols) | H<sub>3</sub>O<sup>+</sup> | Cl<sup>-</sup> | Total |
| 13 | Au+DOX | 236 | 0+50 | 3000 | +50e | 26972 | 72 | 122 | 111796 |
| 14 | Au+An+DOX | 236 | 25+25 | 3000 | 0 | 27288 | 72 | 72 | 112119 |
| 15 | Au+Enol+DOX | 236 | 25+25 | 3000 | +25e | 27269 | 72 | 97 | 112112 |
| 16 | Au+Trans+DOX | 236 | 25+25 | 3000 | +25e | 27288 | 72 | 97 | 112169 |
| 17 | Au+NIR | 236 | 0+50 | 3000 | +50e | 27458 | 72 | 122 | 112054 |
| 18 | Au+An+NIR | 236 | 25+25 | 3000 | 0 | 27469 | 72 | 72 | 112062 |
| 19 | Au+Enol+NIR | 236 | 25+25 | 3000 | +25e | 27458 | 72 | 97 | 112079 |
| 20 | Au+Trans+NIR | 236 | 25+25 | 3000 | +25e | 27490 | 72 | 97 | 112175 |
|  |  | Au | CUR | MTH (mols) | Charge | SOL (mols) | H<sub>3</sub>O<sup>+</sup> | Cl<sup>-</sup> | Total |
| 21 | Au+An | 236 | 2300 | 3000 | -50e | 23818 | 108 | 58 | 92156 |
| 22 | Au+Enol | 236 | 2350 | 3000 | 0 | 23800 | 58 | 58 | 92102 |
| 23 | Au+Trans | 236 | 2350 | 3000 | 0 | 23838 | 58 | 58 | 92216 |
|  | Au+Drug+curcumin on cell membrane | Au | CUR+ Drug (mols) | Membrane | Charge | SOL (mols) | Na<sup>+</sup> | Cl<sup>-</sup> | Total |
| 24 | Au+DOX (Cancer) | 236 | 0+17 | 60913 | -93e | 54106 | 93 | 0 | 224733 |
| 25 | Au+An+DOX (Cancer) | 236 | 25+25 | 60913 | -110e | 53514 | 110 | 0 | 224676 |
| 26 | Au+Enol+DOX (Cancer) | 236 | 25+25 | 60913 | -85e | 53533 | 85 | 0 | 224733 |
| 27 | Au+Trans+DOX (Cancer) | 236 | 25+25 | 60913 | -85e | 53517 | 85 | 0 | 224685 |
| 28 | Au+DOX (DMPC) | 236 | 0+17 | 60416 | +17e | 51741 | 0 | 17 | 217065 |
| 29 | Au+An+DOX (DMPC) | 236 | 25+25 | 60416 | 0e | 51166 | 0 | 0 | 217025 |
| 30 | Au+Enol+DOX (DMPC) | 236 | 25+25 | 60416 | +25e | 51145 | 0 | 25 | 217012 |
| 31 | Au+Trans+DOX (DMPC) | 236 | 25+25 | 60416 | +25e | 51146 | 0 | 25 | 217015 |
| 32 | Au+NIR (Cancer) | 236 | 0+20 | 60913 | -90e | 54195 | 90 | 0 | 224724 |
| 33 | Au+An+NIR (Cancer) | 236 | 25+25 | 60913 | -110e | 53702 | 110 | 0 | 224640 |
| 34 | Au+Enol+NIR (Cancer) | 236 | 25+25 | 60913 | -85e | 53721 | 85 | 0 | 224697 |
| 35 | Au+Trans+NIR (Cancer) | 236 | 25+25 | 60913 | -85e | 53718 | 85 | 0 | 224688 |

All systems subjected to 50,000 steps of steepest-descent energy minimization to remove unfavourable contacts. Subsequently, minimized systems were exposed to a 50 ps NVT equilibration (at temperature 300 K) using 2.0 fs time step. The Berendsen thermostat employed to maintain constant temperature (*71*). This is followed by a 100 ps of NPT equilibration (at 1.0 atm pressure). C-rescale barostat was used to maintain the constant pressure control (*72*). Long-range electrostatics employed particle mesh Ewald (PME) (*73*), while LINCS algorithm constrained hydrogen bonds involving heavy atoms (*74*). The thermodynamically stable configurations were obtained through production NPT MD simulations of 1 μs for Au-curcumin systems and 0.5 μs for the doxorubicin-loaded systems. Potential of mean force (PMF) profiles for Au-curcumin complexes were computed via umbrella sampling, using the inter nanoparticle centre of mass distance as the reaction coordinate (*75*). Windows sampled 20 ns each with 0.1 nm bin size and 1500 kJ/mol/nm^2^ spring constant. For the PMF calculations between drug-loaded Au-nano-complexes and the cancer membrane, the reaction coordinate was defined as the centre-to-centre distance along the membrane normal (z-axis). The umbrella sampling followed by weighted histogram analysis method (WHAM) used to reconstruct the PMF profiles (*76*). Structural analysis included curcumin defining the curcumin region, radial density profiles, surface coverage area (SCA), radius of gyration (Rg), root-mean-square-fluctuations (RMSFs) and tilt angles relative to the Au surface normal providing insights into adsorption behaviour. The simulation methodology detailed previously successfully generated stable trajectories in thermodynamic equilibrium for systems including polymer nanoparticles, proteins, and lipid membranes (*77–82*).

## Results and Analysis

Different forms of curcumin molecules including the negatively charged enolate (An), the neutral enol and trans-keto (Trans) isomers (see the **Figure S1**) as well as their mixtures (An-Enol, Trans-An, and Trans-Enol) were adsorbed onto the surface of 20 Å diameter gold nanoparticle (Au). To explore the formation of thermodynamically stable Au-curcumin complexes, 1.0 µs all-atom molecular dynamics (MD) simulations were performed. The final simulation snapshots of Au-curcumin complexes are shown in **Figures 1a – 1f**: (a) Au+An, (b) Au+Enol, (c) Au+Trans for pure curcumin forms and (d) Au+An-Enol, (e) Au+Trans-An, (f) Au+Trans-Enol) for mixed curcumin cases. The curcumin molecules adsorbed on the Au surface were categorized into four distinct contact-based binding regions, L1 through L4, based on the number of contacts (NOC) each molecule made with the gold nanoparticle (shown in **Figure S2**). Region L1 (blue) consists of curcumin molecules with the highest NOC values (>26, and centre-to-centre distance (r) < 11.7 Å). Regions L2 (red) and L3 (green) correspond to molecules with moderate NOC, ranging from 13 to 26 (11.7 < r < 15.6 Å) and fewer than 13 contacts (15.6 < r < 20.1 Å) respectively. Lastly, region L4 (grey) includes bulk curcumin molecules that do not interact directly with the gold surface (r > 20.1 Å).

**Figure 1:**
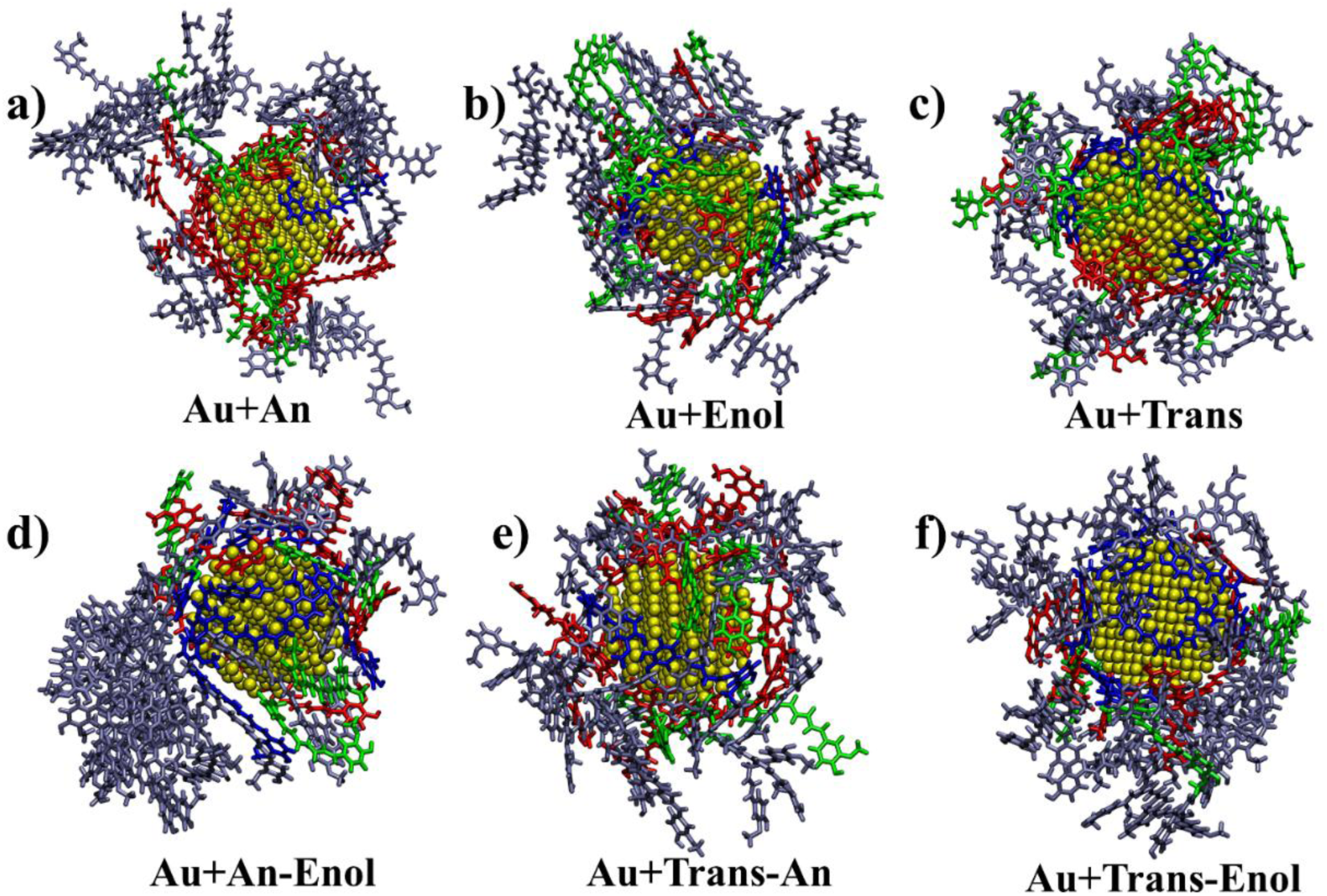
Final simulation snapshots showing pure curcumin molecules (a) Au+An, (b) Au+Enol, (c) Au+Trans and mixed curcumin systems (d) Au+An-Enol, (e) Au+Trans-An, (f) Au+Trans-Enol. The colour of each curcumin molecule corresponds to its contact region with the gold nanoparticle: blue (L1) indicates molecules with the highest number of contacts, red (L2) and green (L3) represent successively fewer contacts, and grey (L4) denotes no contact. Among these complexes, Au+An and Au+An-Enol exhibit the least compact structures, whereas Au+Enol and Au+Trans are the most compact.

The comparison of snapshots shows that curcumin molecules adsorbed in the Au+An complex (shown in **Figure 1a**) tend to protrude outward, which results from unfavourable electrostatic repulsion among negatively charged An species. In contrast, the neutral enol and keto forms in Au+Enol and Au+Trans complexes exhibit a stacked arrangement, providing more effective surface coverage on gold nanoparticles. Additionally, the Au+An complex contains fewer curcumin molecules in region L3 (green) and more in region L4 (grey) compared to the Au+Enol and Au+Trans complexes. For the mixed curcumin complexes, curcumin molecules in the L4 region (grey) of the Au+An-Enol system cluster together through stacking, while in Au+Trans-An and Au+Trans-Enol complexes, these molecules are more evenly distributed across nanoparticle surface. As a result, the Au+Enol and Au+Trans complexes display greater structural compactness, whereas the Au+An and Au+An-Enol complexes form comparatively more loose arrangements.

### 1. Size of the Au – curcumin complex (Rg)

The radius of gyration (Rg) profiles of the Au-curcumin complexes over the simulation time are computed as shown in **Figure S3** and the average values compiled in **Table 2** to compare their overall sizes. The Rg analysis demonstrate that the Au+An (21.9 Å) and Au+An-Enol (20.1 Å) complexes possess larger sizes, whereas the Au+Enol (18.0 Å) and Au+Trans (18.5 Å) complexes show smaller sizes relative to the Au+Trans-An (19.5 Å) and Au+Trans-Enol (19.1 Å) systems. These size discrepancies mainly originate from differences in molecular packing and electrostatic interactions of curcumin molecules. Closer examination of the atomic and molecular arrangements of curcumin molecules on the surface of gold nanoparticle elucidate these differences.

**Table 2:** Radius of gyration and density region values for Au+curcumin complexes and curcumin only, including the changes induced by the presence of the gold nanoparticle.

| Radius of Gyration (Rg) - Å |  |  |  |  |  |  |
| --- | --- | --- | --- | --- | --- | --- |
|  | An | Enol | Trans | An-Enol | Trans-An | Trans-Enol |
| **Curcumin** | 24.5 ± 1.3 | 15.8 ± 0.3 | 17.4 ± 0.7 | 19.4 ± 0.9 | 19.0 ± 1 | 16.7 ± 0.4 |
| **Au + Curcumin** | 21.9 ± 2.1 | 18.0 ± 0.2 | 18.5 ± 0.9 | 20.1 ± 0.8 | 19.5 ± 1.4 | 19.1 ± 0.8 |
| **ΔRg (%)** | -10.5 | 13.9 | 6.1 | 3.8 | 2.6 | 14.1 |
| Thickness of High-Density Region (Tail Region) - Å |  |  |  |  |  |  |
| No. of Peaks (Regions) | 1 (L2)<br>- | 1 (L1 & L2)<br>2 (L4) | 1 (L1 & L2 & L3)<br>2 (L4) | 1 (L1 & L2)<br>2 (L4-Enol) | 1 (L1 & L2)<br>2 (L4-Trans) | 1 (L1 & L2)<br>2 (L4-Enol/Trans) |
| **Curcumin** | 5 (28) | 15 (10) | 14 (13) | 8 (20) | 12 (15) | 14 (12) |
| **Au + Curcumin** | 7 (14) | 8 (8) | 8 (8) | 7 (14) | 7 (10) | 8 (11) |
| **Δ** | 2 (-14) | -7 (-2) | -6 (-5) | -1 (-6) | -5 (-5) | -6 (-1) |

### 2. Atomic configuration inside the Au-curcumin complexes (ρ(r))

Mass density profiles as a function of radial distance from the gold nanoparticle centre were computed for curcumin, ethanol, and water molecules within the Au-curcumin assemblies (**Figure 2**). Furthermore, the widths of the high density and tail regions of the density profiles were tabulated in **Table 2** for comparison.

**Figure 2:**
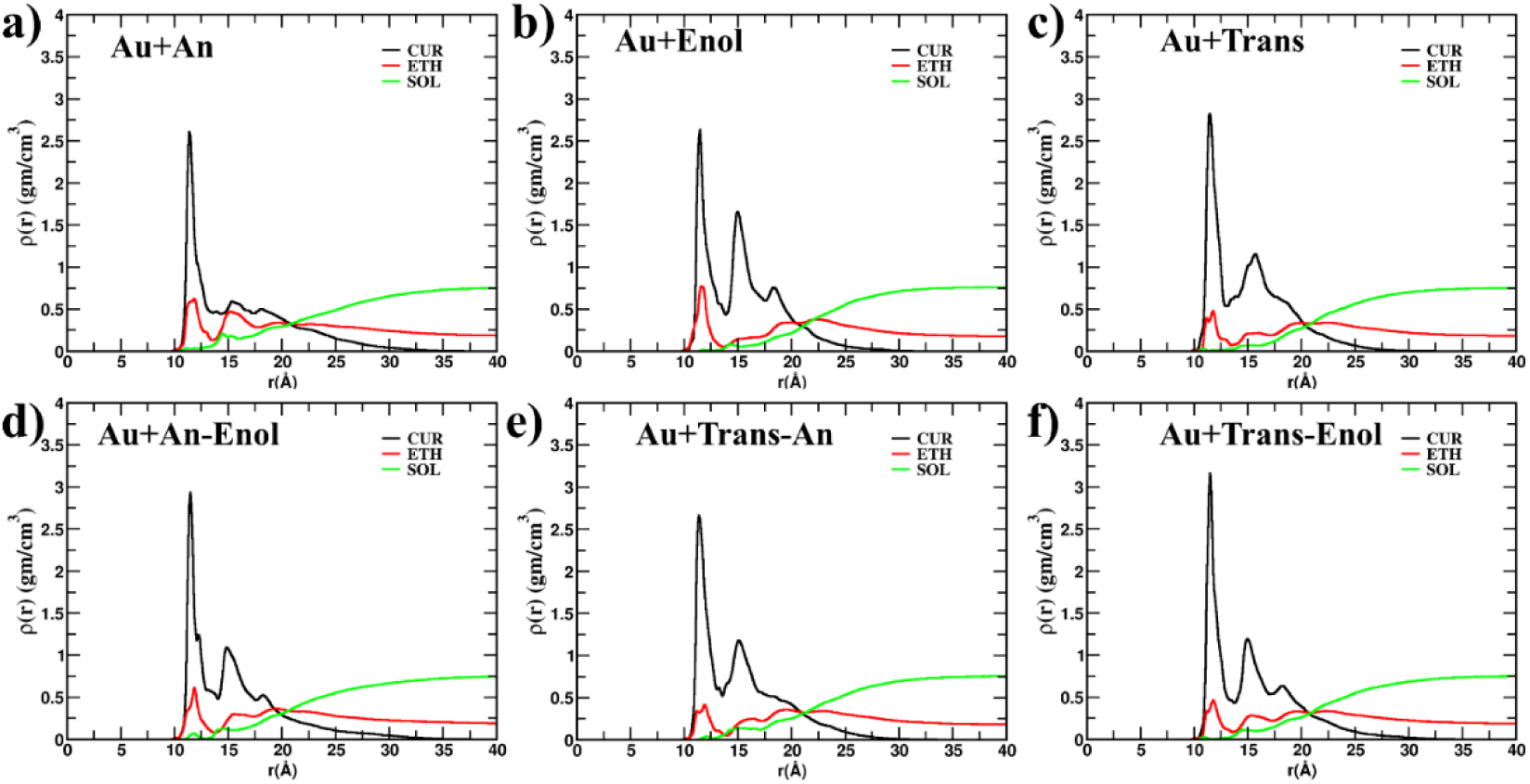
Density profiles of curcumin molecules, ethanol, and water in systems containing (a) Au+An, (b) Au+Enol, (c) Au+Trans, (d) Au+An-Enol, (e) Au+Trans-An, and (f) Au+Trans-Enol, plotted as a function of radial distance from the center of gold nanoparticle. The water density is zero close to the gold nanoparticle surface, while ethanol and curcumin molecules cover the entire nanoparticle surface. All curcumin density profiles display two distinct peaks, except in the Au+An system, which exhibits a single peak. The Au+An and Au+An-Enol systems feature extended tail regions in their density profiles, indicating less compact molecular distributions. In contrast, the density profiles for Au+Trans and Au+Enol systems show shorter tails, reflecting more compact arrangements of curcumin molecules around the nanoparticle.

These profiles indicate that curcumin and ethanol reach their highest density close to the Au-curcumin boundary (∼12 Å), with no detectable water present. This pattern demonstrates complete surface coverage of the gold nanoparticles by curcumin and ethanol, driven by their enhanced van der Waals attraction to the gold surface. All Au-curcumin systems except Au+An display a secondary curcumin density maximum at ∼16 Å, whose magnitude scales with the degree of molecular stacking. The lack of this feature in Au+An arises from the electrostatic repulsion among its anionic curcumin species, which extend outward from the surface. Additionally, the most compact Au+Enol and Au+Trans complexes exhibit sharp tail regions (∼ 8 Å) compared to those of the other complexes (∼ 11-14 Å) (see **Table 2**). To investigate how second-peak and tail region variations in the density profiles relate to curcumin organization, probability distributions of the molecules were calculated for direct comparison.

### 3. Packing of curcumin molecules inside the Au-curcumin complexes

To probe differences in molecular packing, we calculated the probability distribution functions of curcumin molecules in both pure and mixed Au-curcumin systems using a distance cut-off of 7.5 Å, and compared the results (**Figures 3a and 3c**). The analysis shows that, in the anionic Au+An system, configurations with zero or one neighbouring An molecule are most common, whereas in the neutral Au+Enol and Au+Trans systems, arrangements with two or three neighbours occur more frequently. This pattern indicates more pronounced stacking of the charge-neutral Enol and Trans forms in Au+Enol and Au+Trans than for the negatively charged An form in Au+An, in agreement with the missing second peak in the Au+An density profile.

**Figure 3:**
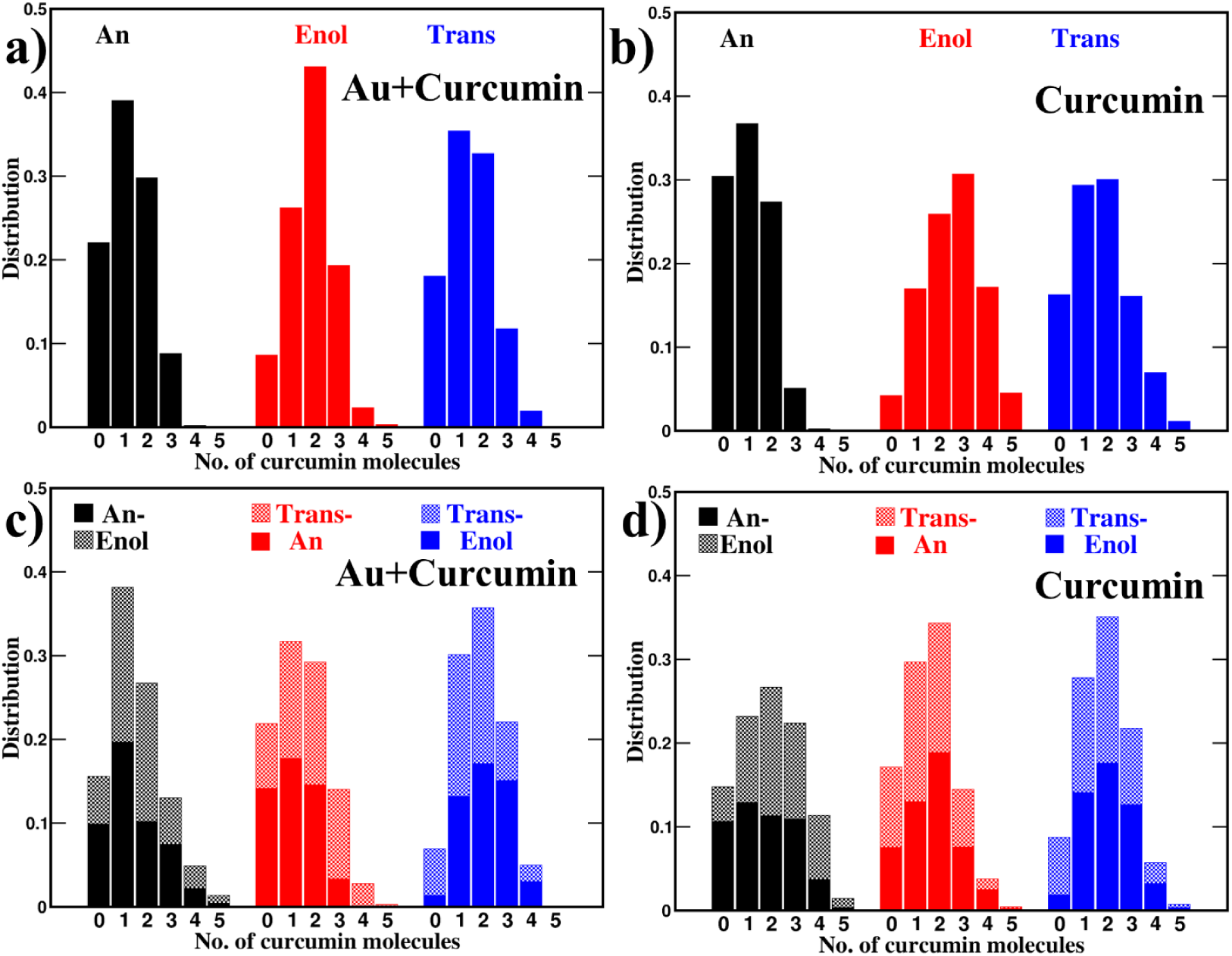
Probability distribution histograms showing the likelihood of finding a given number of curcumin molecules within 7.5Å of another curcumin molecule. Distributions are presented for systems with gold nanoparticles (a and c) and bulk curcumin-only systems (b and d). For gold-containing systems, the probability at zero for Au+An is higher compared to neutral cases (Au+Trans and Au+Enol), indicating reduced aggregation in the anion system. The impact of the gold nanoparticle on the distribution is more pronounced for Enol than for Anion or Trans. In the Au+Trans-Enol mixture, the probability of finding two or three neighbouring curcumin molecules is higher than in other systems, highlighting a more compact and aggregated structure.

In the mixed complexes, each curcumin species largely preserves its intrinsic aggregation behaviour. Systems containing An (Au+An–Enol and Au+Trans–An) still show a greater likelihood of finding zero or one neighbour, consistent with the weaker aggregation seen in pure Au+An. In contrast, Au+Trans–Enol mixture that include the enol form display higher probabilities of having three, and four neighbouring molecules, reflecting stronger stacking and more compact packing driven by enol curcumin. It is in agreement with the high-intensity second-peak and sharp tail region of the density profile.

### 4. Defining regions of curcumin molecules on the gold surface

Curcumin molecules adsorbed on the gold surface were classified into four distinct regions L1, L2, L3 and L4 (**shown in Figure 1**) as described earlier using the number of close contacts (NOC) between curcumin and gold (**Figure S2**). To compare population differences across interfacial regions at the Au-curcumin interface, histograms of the number of curcumin molecules for all four regions are shown in **Figure 4**, with quantitative values in **Table 3**.

**Figure 4:**
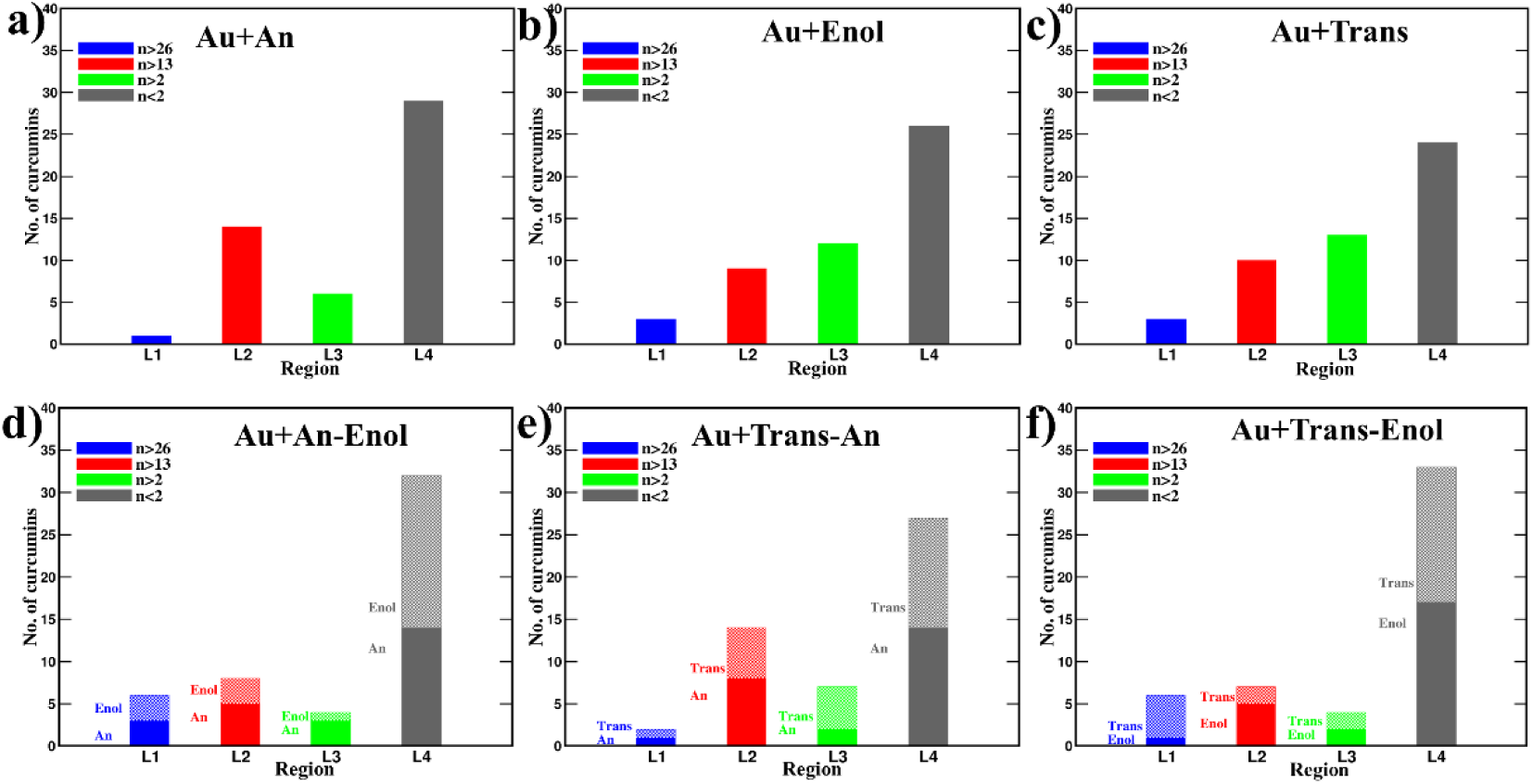
Histograms showing the number of contacts per curcumin molecule for each system: (a) Au+An, (b) Au+Enol, (c) Au+Trans, (d) Au+An-Enol, (e) Au+Trans-An, and (f) Au+Trans-Enol. Curcumin molecules are grouped into four regions (L1–L4) based on their contact counts. The Au+Enol and Au+Trans systems display similar contact distributions. Notably, a higher proportion of anion molecules in Au+An is located in the strongly bound regions (L1 + L2). In mixed systems, the accumulation of anion molecules in L1 + L2 is greater for Au+An-Enol and Au+Trans-An than for their respective Au+Enol and Au+Trans counterparts, indicating a stronger binding preference of anion curcumin in both pure and mixed systems.

**Table 3:** Number of curcumin molecules for each region has been tabulated for Au+curcumin complexes for both pure and mixtures.

| No. of curcumin molecules | L1 | L2 | L3 | L4 |
| --- | --- | --- | --- | --- |
| Au+An | 1 | 14 | 6 | 29 |
| Au+Trans | 3 | 10 | 13 | 24 |
| Au+Enol | 3 | 9 | 12 | 26 |
| Au+An-Enol (An+Enol) | 6 (3+3) | 8 (5+3) | 4 (3+1) | 32 (14+18) |
| Au+Trans-An (Trans+An) | 2 (1+1) | 14 (6+8) | 7 (5+2) | 27 (13+14) |
| Au+Trans-Enol (Trans+Enol) | 6 (5+1) | 7 (2+5) | 4 (2+2) | 33 (16+17) |

In all Au-curcumin complexes, regions L1 and L2 (blue and red) together contain 12–16 strongly bound curcumin molecules (NOC>13). For the Au+An complex, region L2 has the highest population (14 molecules) and L3 the lowest (6 molecules; **Figure 4a**). In neutral Au+Trans and Au+Enol complexes, some curcumin molecules from L2 and L4 migrate to adjacent regions L1 and L3 (**Figures 4b** and **4c**), increasing L1 and L3 populations while decreasing those of L2 and L4 relative to the Au+An complex. These population differences arise primarily from variations in the charge of the curcumin molecules. In mixture systems, An curcumin exhibits stronger binding to gold. In both Au+An-Enol and Au+Trans-An complexes, the inner regions (L1+L2) are predominantly occupied by An curcumin molecules. Conversely, in the Au+Trans-Enol system, the inner regions (L1+L2) are primarily populated by Trans curcumin, underscoring the influence of curcumin type on region population and binding affinity within each complex.

To further investigate the relationship between region populations and their contributions to the overall density profiles, region-by-region density profiles for curcumin were computed (**Figure S4**). In all complexes, the first peaks of the density profiles are predominantly attributed to the L1 and L2 regions, while the second peaks and tail regions mainly arise from the populations in L3 and L4 regions. The packing arrangement of curcumin, its region populations, and the density peaks near the curcumin–gold interface can be directly correlated with the surface area of the gold nanoparticle covered by curcumin molecules in the different gold-curcumin complexes.

### 5. Surface coverage area (SCA) of gold nanoparticle by curcumin molecules

The surface coverage area (SCA) of the gold nanoparticle by curcumin molecules in each region was calculated as a function of time (**Figure S5**), with average percentage values summarized in **Table 4** for comparison. In all systems, the SCA from L1 scales directly with the number of curcumin molecules in this inner region. Regions L1 and L2 together dominate the coverage, accounting for 75-90% of the total due to their extensive contacts with the nanoparticle surface. Region L3, by contrast, contributes only 4-10%, as few atoms from its molecules contact the gold surface directly.

**Table 4:** Surface coverage area (%) of each successive region for Au+curcumin complexes, reported for both pure and mixed systems.

| SCA (%) | L1 | L1 + L2 | L1 + L2 + L3 | L1 + L2 + L3 + L4 | CUR + ETH |
| --- | --- | --- | --- | --- | --- |
| Au+An | 8.2 | 80.3 | 87.4 | 87.9 | 99.9 |
| Au+Enol | 29.0 | 75.1 | 85.7 | 86.8 | 100 |
| Au+Trans | 29.0 | 85.9 | 94.9 | 95.9 | 100 |
| Au+An-Enol | 51.6 | 85.6 | 90.3 | 91.6 | 100 |
| Au+Trans-An | 23.8 | 86.1 | 93.6 | 94.1 | 100 |
| Au+Trans-Enol | 57.8 | 89.8 | 92.2 | 93.4 | 100 |

The Au+Trans complex achieves the highest SCA (96%), surpassing the 87% observed for both Au+An and Au+Enol complexes. The Trans form of curcumin in Au+Trans promotes the most effective anchoring to gold surface, maximizing coverage. In the Enol form, the planar structure favours π-stacking interactions, which reduce SCA. For negatively charged An form, electrostatic repulsions cause the molecules to protrude outward, further lowering coverage. The combined SCA from curcumin and ethanol reaches 100% in all cases, indicating complete exclusion of water from the nanoparticle interface, consistent with the zero water density observed at gold-curcumin interfaces. In mixture systems, SCA values are higher 92% for Au+An-Enol, 94% for Au+Trans-An, and 93% for Au+Trans-Enol demonstrating more effective surface coverage than the pure Au+An or Au+Enol systems.

Although dynamics charge models and quantum calculations can better represent pH-dependent charge states, they are computationally intensive for the time and length scales of the present study. Therefore, fixed-charge models were used, with the corresponding protonation states maintained and H_3_O^+^ and Cl^−^ ions added to ensure charge neutrality and the desired ionic strength. H_3_O^+^ was represented using hydronium-ion parameters as reported in **Table 10**. To examine the effect of counter-ion representation, additional simulations were performed by replacing H_3_O^+^ with Na^+^. As shown in **Figure S6,** the NaCl system exhibits less compact structures and lower surface coverage. However, the overall curcumin-loading mechanism remains similar in both HCl and NaCl conditions, supporting the robustness of the observed mechanism.

### 6. Curcumin size and orientation relative the gold surface as a function of region

To assess the size and orientation of curcumin molecules around the gold nanoparticle, the radius of gyrations (Rg_∥_: parallel and Rg_⊥_: normal to the surface; **Figure S7; Table 5**) and tilt angle (**Figure S8; Table 5**) as function of adsorption regions (L1-L4) were calculated. The radius of gyration (Rg) characterizes the overall three-dimensional extent of a curcumin molecule, while Rg_∥_ and Rg_⊥_, quantify its extent parallel and perpendicular to the nanoparticle surface, respectively. The tilt angle measures the orientation of the curcumin plane relative to the surface normal of the gold nanoparticle. These metrics reveal how Au-curcumin binding influences molecular size and alignment across regions.

**Table 5:** Rg: total, Rg_∥_: parallel and Rg_⊥_: normal to the surface and tilt angle for each region has been tabulated for Au+curcumin complexes for both pure and mixtures.

| Rg (Å) -Total |  |  |  |  |
| --- | --- | --- | --- | --- |
|  | L1 | L2 | L3 | L4 |
| Au+An | 6 ± 0 | 6 ± 0.1 | 6 ± 0.2 | 5.9 ± 0.2 |
| Au+Enol | 5.9 ± 0.4 | 6 ± 0.2 | 5.9 ± 0.2 | 6 ± 0.2 |
| Au+Trans | 5.5 ± 0.5 | 5.7 ± 0.3 | 5.7 ± 0.2 | 5.8 ± 0.2 |
| Au+An-Enol | 5.7 ± 0.2 | 5.9 ± 0.2 | 6 ± 0.1 | 6 ± 0.2 |
| Au+Trans-An | 5.6 ± 0.4 | 5.7 ± 0.4 | 5.6 ± 0.2 | 5.9 ± 0.2 |
| Au+Trans-Enol | 5.2 ± 0.6 | 5.9 ± 0.2 | 5.8 ± 0.4 | 5.9 ± 0.3 |
| Rg (Å) - Parallel |  |  |  |  |
| Au+An | 5.6 ± 0 | 5.2 ± 0.2 | 4.5 ± 0.8 | 5 ± 0.6 |
| Au+Enol | 5.5 ± 0.4 | 5 ± 0.2 | 4.4 ± 1.2 | 5.2 ± 0.7 |
| Au+Trans | 5.3 ± 0.5 | 4.9 ± 0.3 | 4.3 ± 0.7 | 5 ± 0.6 |
| Au+An-Enol | 5.3 ± 0.1 | 5.1 ± 0.6 | 5.2 ± 0.4 | 5.2 ± 0.5 |
| Au+Trans-An | 5.1 ± 0 | 4.8 ± 0.9 | 5.3 ± 0.6 | 4.9 ± 0.7 |
| Au+Trans-Enol | 5.2 ± 0.3 | 4.8 ± 0.7 | 4.6 ± 0.6 | 5 ± 0.7 |
| Rg (Å) - Normal |  |  |  |  |
| Au+An | 1.9 ± 0 | 3 ± 0.5 | 3.7 ± 0.8 | 2.8 ± 0.8 |
| Au+Enol | 1.9 ± 0.3 | 3.3 ± 0.7 | 3.6 ± 1.3 | 2.5 ± 1.1 |
| **Au+Trans** | $1.7 \pm 0.2$ | $2.9 \pm 0.3$ | $3.4 \pm 1$ | $2.5 \pm 0.8$ |
| **Au+An-Enol** | $2.3 \pm 0.7$ | $2.8 \pm 0.83$ | $2.26 \pm 0.49$ | $2.62 \pm 0.73$ |
| **Au+Trans-An** | $2.1 \pm 0.8$ | $3.2 \pm 1.1$ | $2.4 \pm 0.9$ | $2.6 \pm 0.9$ |
| **Au+Trans-Enol** | $2.7 \pm 0.6$ | $3.2 \pm 0.8$ | $3.5 \pm 0.7$ | $2.5 \pm 0.9$ |
| **Tilt angle (°)** |  |  |  |  |
| **Au+An** | $81.9 \pm 0$ | $58.3 \pm 5.9$ | $49.3 \pm 11.6$ | $64.7 \pm 13.6$ |
| **Au+Enol** | $78.1 \pm 6.6$ | $54 \pm 9.4$ | $52.6 \pm 27.3$ | $68.2 \pm 18$ |
| **Au+Trans** | $83.7 \pm 3.8$ | $56.1 \pm 5.3$ | $51.8 \pm 18.1$ | $67.1 \pm 14.7$ |
| **Au+An-Enol** | $78.8 \pm 7.5$ | $62 \pm 4.3$ | $60.7 \pm 20$ | $66.8 \pm 12.6$ |
| **Au+Trans-An** | $88.4 \pm 1.5$ | $53.3 \pm 15.3$ | $50.5 \pm 23.3$ | $69.8 \pm 13$ |
| **Au+Trans-Enol** | $85.8 \pm 5$ | $62.5 \pm 6.6$ | $45.1 \pm 10.6$ | $64.6 \pm 12.9$ |

Trans curcumin exhibits a smaller overall Rg (∼5.6 Å) than the An and Enol forms (∼6.0 Å), attributed to its non-planar geometry. This value remains constant across L1 to L4. In all Au-curcumin complexes, Rg_∥_ (4.5-5.7 Å) is higher than Rg_⊥_ (1.7-3.5 Å), indicating preferential alignment parallel to the surface to maximize contacts. From L1 to L3, Rg_∥_ decreases while Rg_⊥_ increases as Au-curcumin binding weakens and contacts decrease. In unbound L4 region, where curcumin molecules adopt diverse orientations, the opposite trend emerges from L3 to L4.

In connection with this Rg analysis, the tilt angle analysis has been performed. As curcumin molecules in L1 align more parallel to the surface, their tilt angles (75°-90°) relative to the surface normal (**Figure S9a**) are higher compared to L2 (40°-70°) and L3 (20°-80°) (**Figure S9 b-c**). The alignment decreases from L1 to L3, and the tilt angle distributions become broader with lower mean values. However, the Au+Enol system shows a wider spread of tilt angles due to a higher degree of stacking compared to other Au-curcumin complexes. Both Rg and tilt angle analyses indicate a reduction in curcumin alignment to gold surface in the mixed Au-curcumin complexes. Moreover, the larger error bars in the tilt angles of the L4 region suggest that curcumin molecules in the bulk are more flexible than those in the inner curcumin regions of the complexes. To further understand these changes in relation to curcumin flexibility, root-mean-square fluctuations (RMSF) analysis of curcumin molecules was performed.

### 7. Flexibility of curcumin regions in Au+curcumin complex (RMSF)

Probing the changes in the flexibility of curcumin regions in Au-curcumin complexes, the root-mean-square fluctuations (RMSF) were computed as a function of region, as shown in **Figure 5** for comparison. The RMSF profiles indicate that flexibility increases from L1 to L4, reflecting reduced binding to the gold nanoparticle across all complexes. Among the complexes, curcumin in the Au+Trans system exhibits the highest flexibility, whereas the lowest flexibility is observed in the Au+Enol complex. This difference arises because the non-planar geometry of Trans curcumin leads to lower packing efficiency, while the planar Enol form favours strong stacking interactions. In the Au+An complex, the negatively charged An curcumins protrude outward to minimize electrostatic repulsions, resulting in moderate flexibility. In mixed Au-curcumin complexes, curcumin flexibility is similar, as repulsions between An and stacking interactions between Enol are reduced. Overall, this interplay of molecular orientation, packing density, and binding affinity governs the flexibility differences observed among curcumin forms and regions.

**Figure 5:**
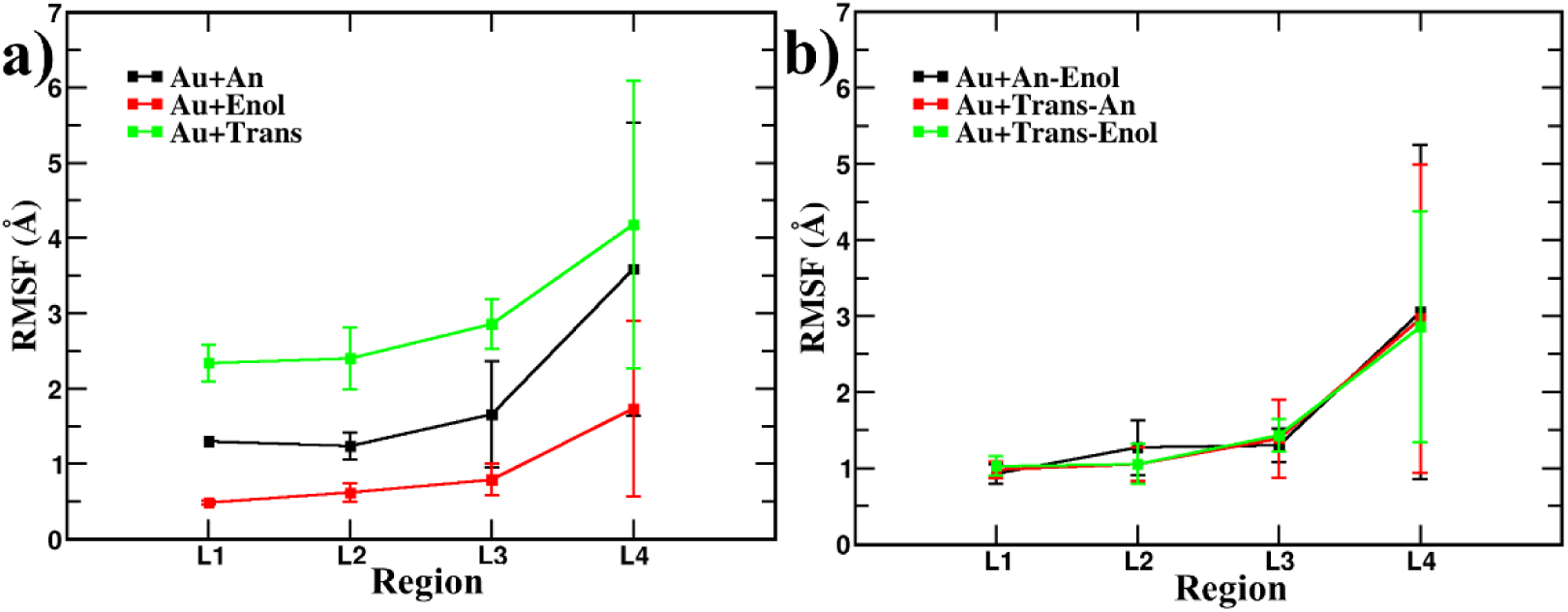
Root mean square fluctuations (RMSF) of curcumin molecules in the presence of a gold nanoparticle, plotted as a function of contact region: (a) pure curcumin systems (Au+An, Au+Enol, Au+Trans) and (b) mixture systems (Au+An-Enol, Au+Trans-An, Au+Trans-Enol). RMSF increases progressively from the innermost to outermost region (L1 to L4), indicating that L1 molecules are the least flexible while L4 molecules exhibit the highest flexibility. Among pure systems, Au+Trans displays the greatest flexibility, whereas Au+Enol shows the lowest. In all mixture systems, RMSF values are similar for all systems, suggesting comparable flexibility among the constituent curcumin types.

### 8. Structural impact of Au-curcumin complexation relative to curcumin in water

Au-curcumin interactions alter molecular organization, as revealed by comparing curcumin structures in gold nanoparticle complexes versus free clusters in aqueous solution (**Figures 3** and **S3**). Snapshots of curcumin aggregates lacking the gold nanoparticle appear in **Figure 6**, with key metrics summarized in **Tables 2** and **6**. In water, neutral Enol, Trans and Trans-Enol curcumin aggregates display high compactness form the extensive π-π stacking. Negatively charged An curcumin aggregates show the least compactness, while mixed An-Enol and Trans-An forms fall in between, primarily due to charge induced electrostatic repulsion.

**Figure 6:**
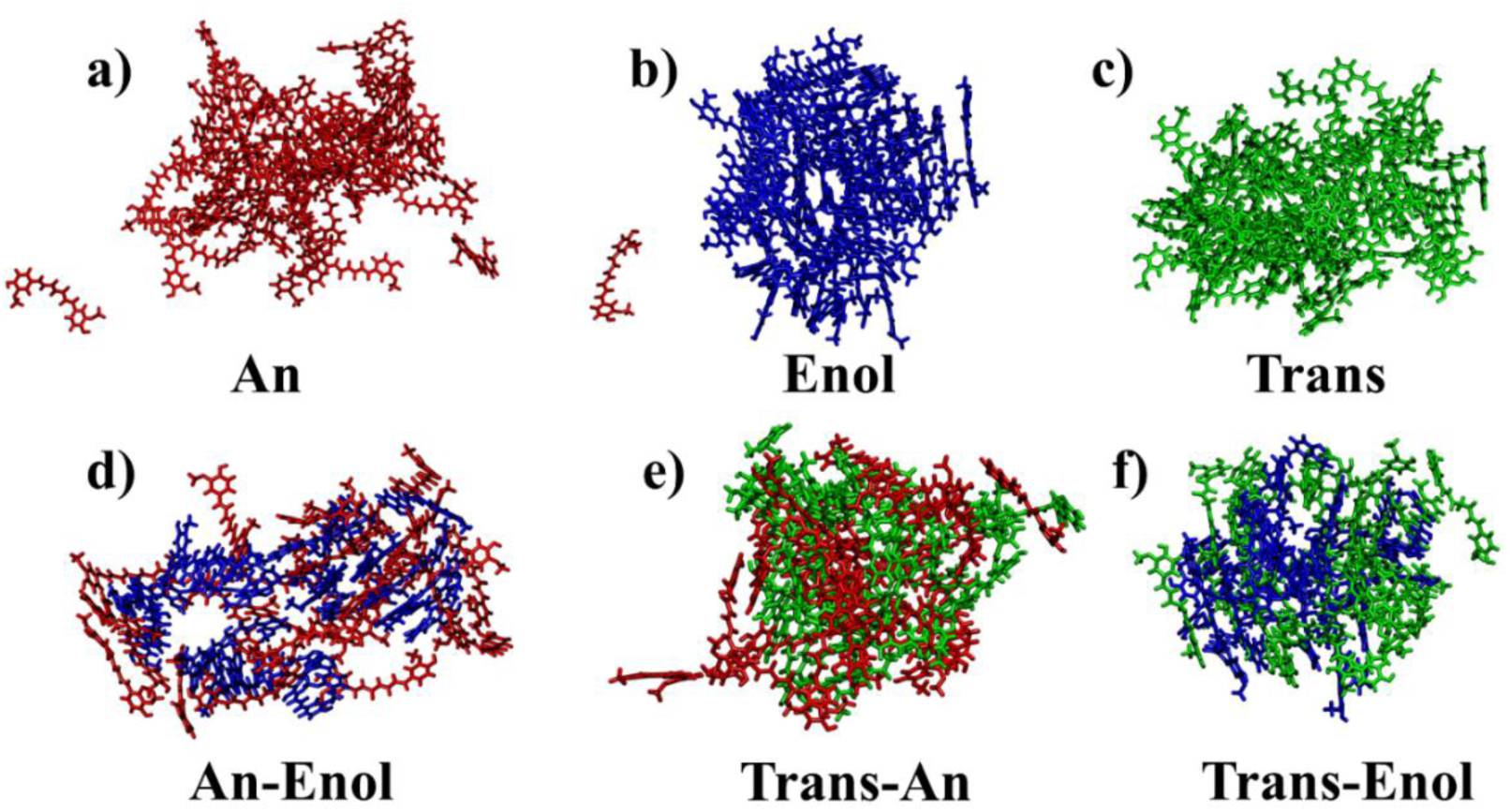
Final simulation snapshots of pure curcumin molecules (a) An, (b) Enol, and (c) Trans and mixture systems (d) An-Enol, (e) Trans-An, and (f) Trans-Enol in the absence of a gold nanoparticle. The An system exhibits poor aggregation and loose packing, attributed to electrostatic repulsion between negatively charged curcumin molecules. In contrast, neutral systems (Trans, Enol, and Trans-Enol) display the most compact molecular packing. Semi-neutral mixtures (Trans-An and An-Enol) are less compact than the fully neutral cases, indicating the influence of charge on aggregation behaviour.

**Table 6:** Curcumin probability (%) is tabulated for both Au+curcumin and curcumin only, including the changes induced by the presence of the gold nanoparticle.

| Probability (%) | 0 | 1 | 2 | 3 | 4 | 5 | Δ0 | Δ1 | Δ2 | Δ3 | Δ4 | Δ5 |
| --- | --- | --- | --- | --- | --- | --- | --- | --- | --- | --- | --- | --- |
| **An** | 31 | 37 | 27 | 5 | 0 | 0 | -- | -- | -- | -- | -- | -- |
| **Au+An** | 22 | 39 | 30 | 9 | 0 | 0 | -9 | 2 | 3 | 4 | 0 | 0 |
| **Enol** | 4 | 17 | 26 | 31 | 17 | 5 | -- | -- | -- | -- | -- | -- |
| **Au+Enol** | 9 | 26 | 43 | 20 | 2 | 0 | 5 | 9 | 17 | -11 | -15 | -5 |
| **Trans** | 16 | 30 | 30 | 16 | 7 | 1 | -- | -- | -- | -- | -- | -- |
| **Au+Trans** | 18 | 35 | 33 | 12 | 2 | 0 | 2 | 5 | 3 | -4 | -5 | -1 |
| **An-Enol** | 15 | 23 | 27 | 22 | 11 | 2 | -- | -- | -- | -- | -- | -- |
| **Au+An-Enol** | 16 | 38 | 27 | 13 | 5 | 1 | 1 | 15 | 0 | -9 | -6 | -1 |
| **Trans-An** | 17 | 30 | 34 | 15 | 4 | 0 | -- | -- | -- | -- | -- | -- |
| **Au+Trans-An** | 22 | 32 | 29 | 14 | 3 | 0 | 5 | 2 | -5 | -1 | -1 | 0 |
| **Trans-Enol** | 8 | 28 | 35 | 22 | 6 | 1 | -- | -- | -- | -- | -- | -- |
| **Au+Trans-Enol** | 7 | 30 | 36 | 22 | 5 | 0 | -1 | 2 | 1 | 0 | -1 | -1 |

Binding to the gold nanoparticle compacts An curcumin aggregates more than their free form in water, as gold nanoparticle surface interaction outweigh intermolecular repulsions. This yields a smaller radius of gyration (Rg; Δ = −10.5 Å), broader high-density regions (Δwidth = +2 Å), and narrower tail regions (Δwidth = −14 Å) in the density profiles (**Figure S10**). For the initially compact neutral Enol, Trans and Trans-Enol aggregates, complexation expands their overall size (larger Rg in **Table 2** relative to free clusters) and further narrows high-density regions.

Furthermore, the stacking patterns were analysed by comparing distribution of curcumin molecules with 0-5 neighbouring curcumins in Au-curcumin complexes versus free aqueous aggregates (**Table 6**). For the less compact An aggregates, complexation enhances higher order stacking, with 9% of isolated curcumins in water forming more extensive stacks upon gold nanoparticle binding. In contrasts, the highly compact Enol, Trans, and An-Enol aggregates reduce their stacking extent upon complexation, redistributing molecules to concentrate compactness near the Au-curcumin interface. This shift increases the fraction of molecules with only 1-2 neighbours compared to their free aqueous forms.

The number of water and ethanol molecules associated with the curcumin complexes within 5 Å cut-off are computed and presented for comparison in **Table 7** to evaluate variations in solvation. These counts peaked for the anionic enolate (An) curcumin and were minimal for the Enol curcumin. Gold nanoparticle binding decreased solvation in An case (25% lesser water molecules and 5% lesser ethanol molecules) compared to free curcumin clusters in water, whereas it enhanced solvation in neutral Au+Enol (+5% water, +39% ethanol) and Au+Trans (+12% water, +13% ethanol) systems. These trends are due to increased structural compactness in Enolate (An) and decreased stacking interactions in Enol and Trans forms after Au binding. Greater solvation in the neutral Au+Enol and Au+Trans complexes suggests superior bioavailability relative to unbound curcumin.

**Table 7:** Number of water molecules and ethanol molecules is tabulated for both Au+curcumin and curcumin only.

| **Number of Bound Water Molecules** |  |  |  |  |  |  |
| --- | --- | --- | --- | --- | --- | --- |
|  | **An** | **Enol** | **Trans** | **An-Enol** | **Trans-An** | **Trans-Enol** |
| **Curcumin** | $786 \pm 46.1$ | $385.6 \pm 30.3$ | $500 \pm 48.3$ | $490.8 \pm 37.5$ | $503.4 \pm 31.7$ | $451.4 \pm 31.1$ |
| **Au + Curcumin** | $591.3 \pm 32.0$ | $404.3 \pm 23.3$ | $562.1 \pm 38.1$ | $489.9 \pm 33.9$ | $499.6 \pm 35.4$ | $451.2 \pm 34$ |
| **Δ (%)** | -25 | 5 | 12 | 0 | -1 | 0 |
| **Number of Bound Ethanol Molecules** |  |  |  |  |  |  |
|  | **An** | **Enol** | **Trans** | **An-Enol** | **Trans-An** | **Trans-Enol** |
| **Curcumin** | $345.7 \pm 19.5$ | $172.2 \pm 10.6$ | $223.7 \pm 14.9$ | $229.4 \pm 15.1$ | $212.2 \pm 11.8$ | $206 \pm 12.2$ |
| **Au + Curcumin** | $329.5 \pm 15.3$ | $239 \pm 11.3$ | $252.3 \pm 12.6$ | $266.9 \pm 12.6$ | $264.1 \pm 10.3$ | $248.3 \pm 12.7$ |
| **Δ (%)** | -5 | 39 | 13 | 16 | 24 | 21 |

### 9. Influence of ethanol-water and methanol-water solvents on the structure Au+curcumin complexes

The influence of solvent on the Au+curcumin complexes was examined by substituting ethanol with methanol. Final snapshots of pure curcumin complexes in methanol are shown in **Figure S11**, and key comparative properties such as aggregate size (**Figure S12**), atomic distribution (**Figure S13**), and surface coverage area (SCA), are summarized in **Table 8**. Analysis of the sizes together with simulation snapshots indicates that the complexes are more compact in methanol, reflecting stronger Au-curcumin binding than in ethanol. Consistently, the density profiles of the complexes exhibit higher-intensity peaks in methanol compared to ethanol (**Figure S13**). Moreover, the SCA values of the gold nanoparticle in the An, enol, and trans curcumin complexes increase by 9%, 12%, and 4%, respectively in methanol relative ethanol. This behaviour arises because ethanol better stabilises the more extended An curcumin aggregates and Enol stacks in the complexes than methanol does (**Figure S14**), leading to lower SCA values for gold nanoparticle in ethanol.

**Table 8:** Radius of gyration, density thickness, and surface coverage (SCA %) of pure curcumin complexes in ethanol and methanol, are compared.

| Radius of Gyration (Rg) - Å |  |  |  |
| --- | --- | --- | --- |
|  | Au+An | Au+Enol | Au+Trans |
| Au + Curcumin (ETH) | 21.9 ± 2.1 | 18.0 ± 0.2 | 18.5 ± 0.9 |
| Au + Curcumin (MTH) | 19.0 ± 0.4 | 16.8 ± 0.1 | 17.2 ± 0.2 |
| Thickness of High Density Region (Tail Region) - Å |  |  |  |
| Au + Curcumin (ETH) | 7 (14) | 8 (8) | 8 (8) |
| Au + Curcumin (MTH) | 7 (13) | 8 (7) | 8 (7) |
| Total CUR SCA (%) |  |  |  |
| Au + Curcumin (ETH) | 87.9 | 86.8 | 95.9 |
| Au + Curcumin (MTH) | 96.9 | 98.5 | 100.0 |

**Table 9:** Details of the lipid and cholesterol composition in modelled cancer cell membrane.

| No. of Lipids | POPS | CHL | PSM | POPC | POPE |
| --- | --- | --- | --- | --- | --- |
| Upper leaflet | 46 | 43 | 76 | 69 | 22 |
| Lower leaflet | 64 | 43 | 9 | 32 | 108 |
| Total Number | 110 | 86 | 85 | 101 | 130 |

**Table 10:** Details of the vdw parameters (σ & ε) for the Ions and gold atoms used in our study.

| Atom | $\sigma$ (nm) | $\epsilon$ (kJ/mol) | Charge (e) | Ref |
| --- | --- | --- | --- | --- |
| H<sub>3</sub>O<sup>+</sup> | 0.238 | 0.033 | +1 | (69) |
| Na<sup>+</sup> | 0.244 | 0.036 | +1 | (91) |
| Cl<sup>-</sup> | 0.448 | 0.149 | -1 | (91) |
| Au | 0.263 | 22.133 | 0 | (61) |
| Au* | 0.263 | 43.06 | 0 | (83) |

To elucidate the origin of these differences in stabilization, the number of bound ethanol and methanol molecules, as well as the close-contacts between ethanol/methanol molecules and curcumin, were compared for the corresponding Au-curcumin complexes, as shown in **Figure S15**. The comparative analysis reveals that a greater number of ethanol molecules are bound to the Au-curcumin complexes than methanol molecules. This enhanced ethanol solvation is accompanied by a greater number of ethanol-curcumin close contacts, indicating a stronger association of ethanol with curcumin relative to methanol. This enhanced ethanol solvation may be attributed to the greater hydrophobicity of ethanol compared with methanol, which may promote closer association with the hydrophobic moieties of curcumin and consequently contribute to improved solvation and stabilization of the complex.

The distribution of curcumin molecules across different regions was also compared between the two solvents, as shown in **Figure S16**. For all complexes, the number of curcumin molecules in the innermost regions (L1 + L2) is higher in methanol than in ethanol. This enhanced binding in methanol reduces the conformational flexibility of the adsorbed curcumin regions on the gold surface compared to ethanol (see **Figure S17**). Among the three curcumin forms, the solvent effect is most pronounced for the An curcumin, whereas Trans form shows the least sensitivity to solvent variation. Overall, these results highlight the critical role of the solvent environment in governing curcumin binding and spatial organization at the Au-curcumin interface.

### 10. Binding free energies and effective interactions of Au-curcumin complexes: PMF (potential of mean force) calculations

To assess the stability differences among the Au-curcumin complexes, the total binding free energies and their pairwise contributions (Au-curcumin, Au-ethanol, and curcumin-ethanol) were computed using MMPBSA (Molecular Mechanics Poisson–Boltzmann Surface Area) method (**Figure S18**). These binding energies are summarised in **Table S3**. The analysis reveals that the Au-An curcumin complex exhibits the highest stability, followed by intermediate stability in the Au+Enol, Au+ Trans, and Au+Trans-Enol complexes show the lower stability. These variations arise primarily from differences in the Au-curcumin and curcumin-ethanol interactions. Additionally, potential of mean force (PMF) calculations were performed on various Au-curcumin complexes to quantify their effective interactions. These profiles were compared with that of bare gold nanoparticles (**Figure 7**). The An curcumin fully screens Au-Au interactions (ΔG = −20 kcal/mol, 100% efficient), whereas Enol and Trans curcumin achieve only 50% screening (ΔG = −10 kcal/mol). Among mixed complexes, the An-Enol curcumin provides the strongest screening (ΔG = −5 kcal/mol, 75% efficient), while Trans-An curcumin shows the weakest. Moreover, the effective interaction range in Au-curcumin complexes doubles to 20 Å (minimum at ξ = 37 Å) compared to 10 Å (minimum at ξ = 27 Å) for bare gold nanoparticles. These effects arise from electrostatic repulsions by negatively charged curcumin, π-stacking at contact interfaces, and stabilization of the aqueous ethanol solvent.

**Figure 7:**
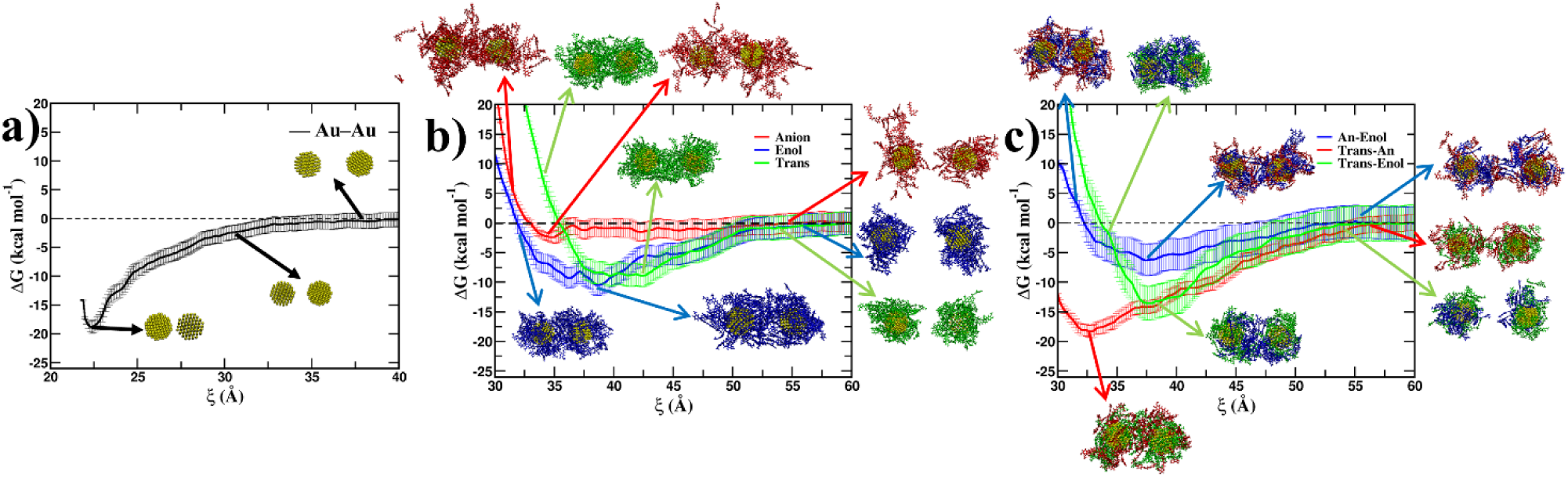
The PMF profiles are plotted for (a) only gold nanoparticle (b) pure curcumin cases (anion, trans and enol) and (c) mixtures (an-enol, trans-an and trans-enol). An curcumin completely shields Au-Au interactions, proving most effective at screening, while Enol and Trans forms provide only partial coverage. Among mixtures, An-Enol curcumin offers the strongest screening, with Trans-An showing the weakest performance.

### 11. Doxorubicin (DOX) loading onto Au-curcumin complexes

Au-curcumin-DOX complexes were constructed to investigate variations in doxorubicin (DOX) loading in the presence of different curcumin forms (An, Enol and Trans) on Au nanoparticles, in comparison with DOX alone under ethanol solvent conditions. The final simulation snapshots are shown in **Figure S19**. In the absence of curcumin, 18 out of 50 DOX molecules (∼38%) were adsorbed onto the Au nanoparticle surface, exhibiting a radius of gyration (Rg) of 16.1 Å, despite the net positive charge of DOX. In contrast, in the presence of the curcumin, the curcumin molecules preferentially occupy the first coordination layer on the Au nanoparticle, while DOX molecules from stacked complexes with all curcumin forms, effectively screening their electrostatic repulsions. For the negatively charged An, the stacking is well-organized and adopts a more expanded conformation (Rg = 21.1 Å), whereas more compact structures (Rg = 18.8 Å) are observed for the neutral Enol and Trans forms. Overall, curcumin-mediated complexation significantly enhances DOX loading onto the Au nanoparticle, increasing it from 38% to approximately 85-98% with a surface coverage of 80-85%. Furthermore, MMPBSA calculations indicate that the binding of curcumin and DOX to the Au nanoparticle surface is primarily driven by favourable van-der-Waal (vdW) interactions.

The Au-DOX-curcumin complexes were further equilibrated for 50 ns to assess the impact of the absence of ethanol on their structural properties. The corresponding simulation snapshots and structural analysis are presented in **Figure 8**. In the absence of ethanol, the Au+DOX complex exhibits minimal structural changes, whereas curcumin adsorption significantly increases to 90-98%, resulting almost complete surface coverage of Au nanoparticle. Furthermore, neutral complexes involving the Enol and Trans forms adopt more compact and mechanically stable configurations, driven by enhanced vdW interactions governing Au-curcumin/DOX binding (**Table S5, Figure S20**). These equilibrated structures were subsequently utilized to investigate the interactions of Au-DOX-curcumin complexes with cancer cell membranes. Furthermore, the robustness of DOX-loading onto the Au surface was examined by varying key simulation parameters, including the initial number of DOX molecules (30, 40 and 50), box size, initial placement, and simulation duration. The alternative Au* force field reported by Jacobson et al (*83*). was also evaluated and compared with the original Au force field of Heinz et al. (*61*). These analyses, shown in **Figure S21**, demonstrate that variations in the Au force-field parameters and other initial MD conditions have minimal influence on the DOX-loading mechanism, both in presence and absence of curcumin. The interaction energies of Au-DOX and Au-curcumin complexes containing An, Enol, and Trans forms were obtained from MD trajectories and compared with van-der-Waal energies coming from MM-PBSA calculations in the **Tables S4 & S6**. The comparison of the interaction energies further confirms that the neural curcumin forms (Enol and Trans) exhibit stronger binding to the Au surface than the anionic (An) curcumin from and DOX.

**Figure 8:**
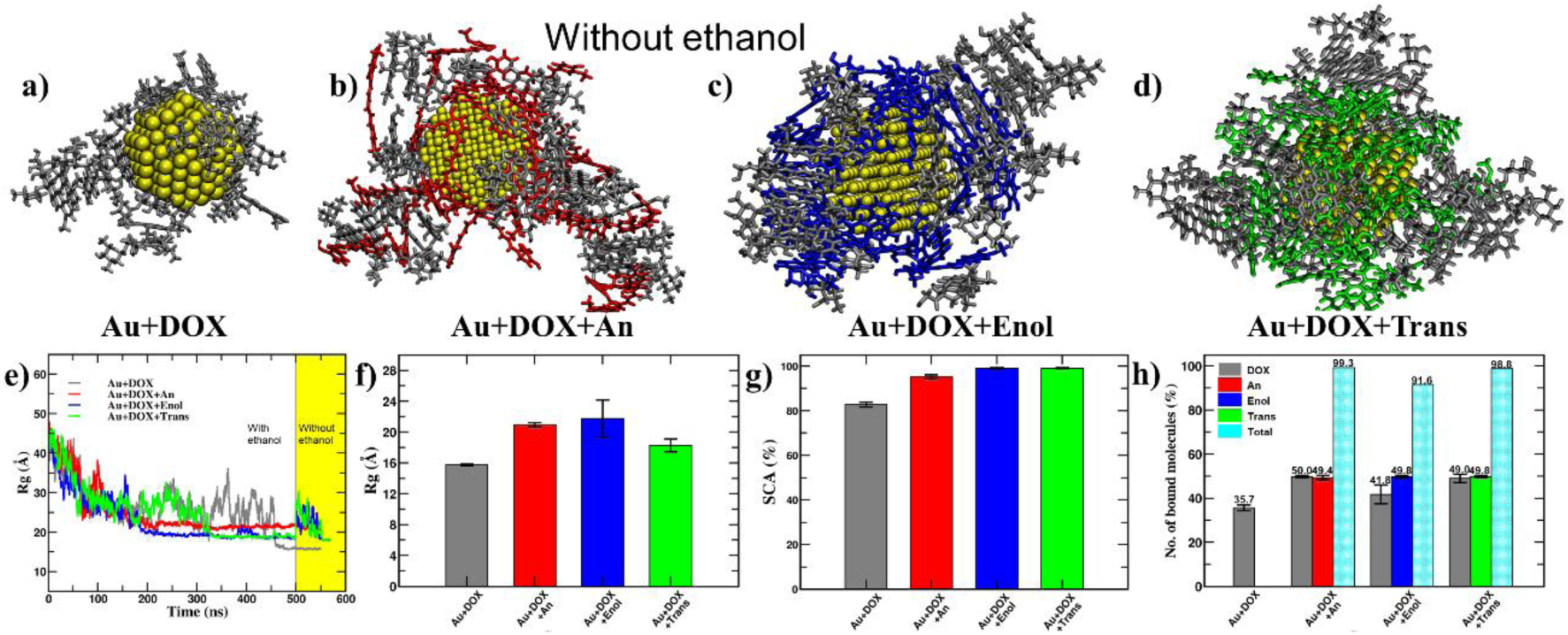
Molecular arrangement of curcumin (An, Enol, and Trans) and doxorubicin (DOX) in final snapshots of (a) Au+DOX, (b) Au+DOX+An, (c) Au+DOX+Enol, and (d) Au+DOX+Trans, complexes in the absence of ethanol. Only 17 out of 50 DOX molecules adsorb to the Au nanoparticle in Au+DOX, whereas curcumin enhances DOX loading and nanocomposite stability. This is supported by the radius of gyration (e & f), % surface coverage area (g), and number of bound molecules (h) to Au nanoparticle, indicating increased DOX binding in the presence of curcumin.

To assess the stability of DOX and curcumin molecules within the Au+DOX+curcumin complexes, their root-mean-square fluctuations (RMSF), intermolecular contacts distributions, and mean-square displacements (MSD) were analyzed and compared, shown in **Figure S22**. The comparative RMSD and MSD analyses reveal that the DOX and curcumin molecules in the Au+DOX+Enol and Au+DOX+Trans complexes exhibit reduced conformational flexibility and molecular mobility compared with those in the Au+DOX+An complex. These results suggest that the neutral Enol and Trans forms of curcumin confer greater stability to complexes, which may be particularly relevant under tumor microenvironment conditions. In contrast, the electrostatic repulsion between positively charged DOX molecules is effectively screened by the negatively charged An curcumin molecules, resulting in an increased number of close intermolecular contacts and enhanced molecular flexibility. Furthermore, PMF profiles of DOX as a function of its radial distance from the Au nanoparticle indicate slower DOX release from the complexes containing neutral Enol and Trans forms, consistent with their higher structural stability, shown in **Figure S22**. The PMF and MSD analyses suggest that the characteristic timescale associated with DOX release is substantially longer than the timescale accessible within MD simulations. However, the actual tumour microenvironment is considerably more complex and includes negatively charged lipids, cholesterol, and membrane-associated proteins, and other components that substantially influence nanoparticle-membrane interactions and drug-release behaviour.

### 12. Interaction of Au+DOX+curcumin complexes with cancer cell membrane

To investigate the binding of the Au+DOX+curcumin complexes to negatively charged cancer cell membrane and zwitterionic DMPC lipid membranes, the complexes were initially positioned near the lipid membrane surface and subjected to 500 ns MD simulations to obtain stable conformations. The final simulation snapshots, and corresponding analyses are shown in **Figure 9**. The centre to centre distance (Δz) profiles as a function of simulation time indicate that the complexes reach a stable membrane-bound state after approximately 100 ns. The net positively charged Au+DOX complex adopts a relatively compact conformation, characterized by a smaller radius of gyration (Rg=16.1 Å) compared with those of the Au+DOX+curcumin complexes. This compact structure facilitates favourable electrostatic interactions with the negatively charged cancer cell membrane, resulting in the shortest centre to centre distance (Δz = 32 Å) among all the investigated systems. In comparison, the Au+DOX+An and Au+DOX+Trans complexes exhibit larger center-to-center distances (Δz = 40 Å), whereas the Au+DOX+Enol complex exhibits an intermediate distance (Δz = 35 Å). However, these complexes exhibit relatively slower adsorption onto DMPC membrane, occurring over 300 – 500 ns, and maintain larger separation ((Δz ∼ 42 Å).

**Figure 9:**
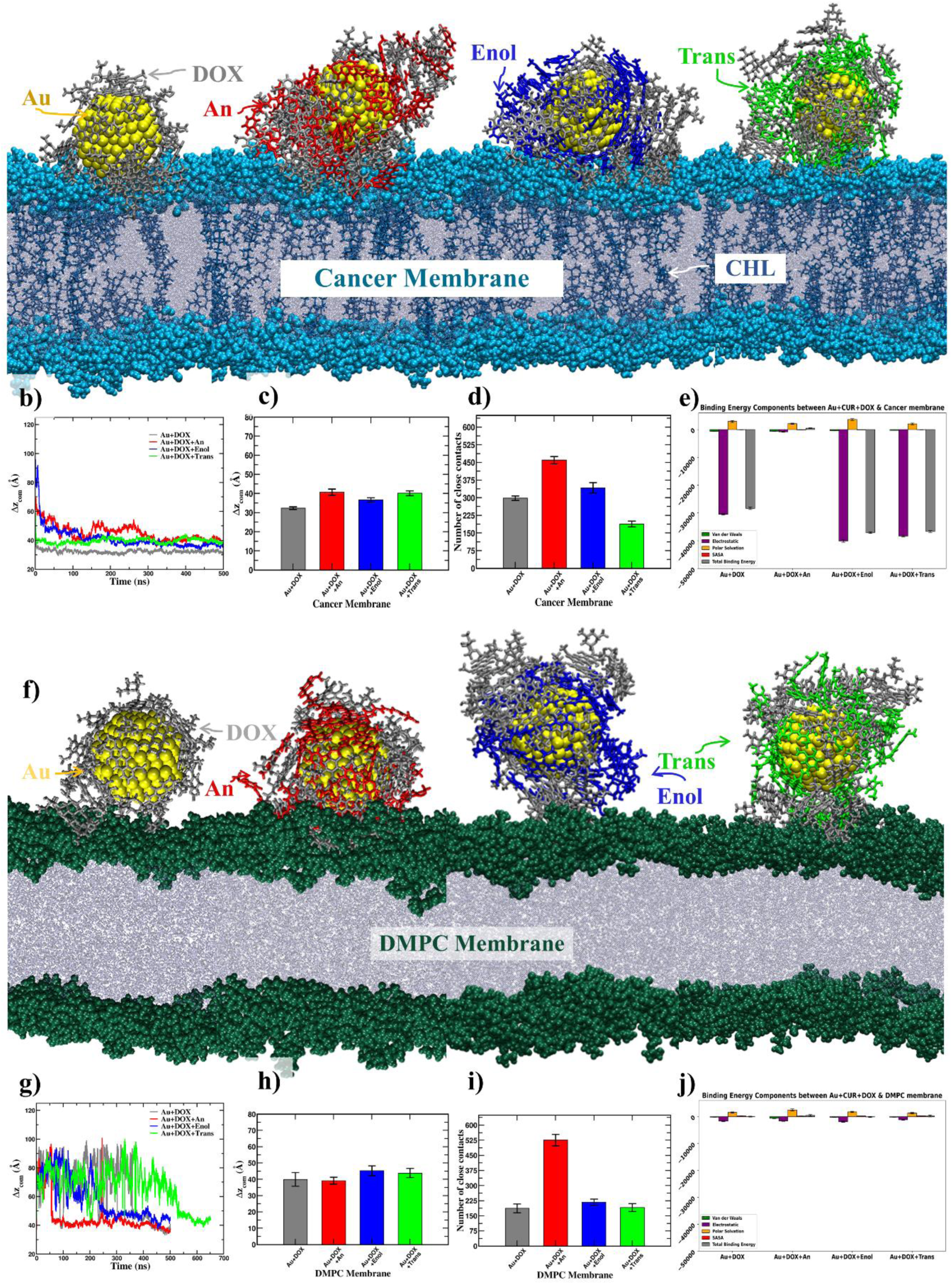
Binding of (a) Au+DOX, Au+DOX+An, Au+DOX+Enol, and Au+DOX+Trans nanocomposites to the cancer cell membrane and (f) DMPC membrane after 0.5 μs production MD. The complexes containing neutral curcumin (Au+DOX+Enol and Au+DOX+Trans) exhibit stronger binding to the cancer cell membrane, whereas Au+DOX+An shows weaker interactions compared with Au+DOX. These trends are consistent with the (b, c, g & h) centre-to-centre distances, (d & i) number of close-contacts and (e & j) MMPBSA analyses. In contrast, the Au+DOX+Curcumin complexes exhibit weaker interactions with zwitterionic DMPC lipid membrane than with the cancer cell membrane, indicating a preferential interaction with the cancer cell membrane.

Furthermore, comparison of the center-to-center distance profiles along with close-contact analyses demonstrates that the Au+DOX+curcumin complexes interact more strongly with the negatively charged cancer cell membrane than with the zwitterionic DMPC membrane. This enhanced membrane association can primarily be attributed to favourable electrostatic interactions between the positively charged DOX moiety and the negatively charged lipid head-groups of the cancer cell membrane. These differences are particularly pronounced for the Au+DOX, Au+DOX+Enol, and Au+DOX+Trans systems compared with the Au+DOX+An complex. The weaker interaction observed for Au+DOX+An can be attributed to electrostatic charge screening by the negatively charged An form of curcumin, which partially neutralizes the positive charge of DOX and consequently reduces the net positive charge of the Au+DOX+An complex. This reduction in net positive charge weakens the electrostatic attraction between the complex and the negatively charged cancer cell membrane.

MM-PBSA is a widely used approach for estimating protein-ligand (*84,85*) and protein-protein (*77*) binding free energies. Although MM-PBSA calculations can overestimate electrostatic and polar solvation contributions in metallic nanoparticle systems because of the use of continuum implicit-solvent models (*86*), the method can nevertheless provide useful insights into relative differences in binding affinity. Accordingly, comparison of the MM-PBSA binding free energies further reveals that complexation of positively charged DOX with the neutral Au-curcumin systems (Enol and Trans) enhances membrane binding relative to the Au+DOX complex. This enhanced binding may arise from the spreading of DOX molecules over the stacked curcumin structures, which increases the effective interaction area between the positively charged DOX moieties and the negatively charged lipid head groups of the cancer cell membrane, thereby strengthening electrostatic attraction. Despite this enhanced interaction with the negatively charged cancer cell membrane, the Au+DOX+Enol, and Au+DOX+Trans complexes exhibit comparatively weaker binding to the zwitterionic DMPC membrane, consistent with the overall charge neutrality of later. In contrast, stacking of negatively charged An curcumin with DOX partially screens the electrostatic repulsions between DOX moieties, resulting in the lowest binding affinity among the investigated complexes for both the cancer cell and DMPC membranes. Overall, these findings indicate that incorporation of curcumin into Au+DOX composites modulates their membrane binding behaviour and can enhance their preferential association with negatively charged cancer cell membranes. In particular, neutral curcumin forms (Enol and Trans) promote stronger membrane binding, whereas the negatively charged An form attenuated the electrostatic interaction through charge screening. These results provide molecular-level insight into how the protonation state of curcumin influences the membrane interactions and biocompatibility of Au+DOX+curcumin nanocomposites.

### 13. Free energy of binding of Au+DOX+curcumin complexes to the cancer cell membrane

In addition to the MM-PBSA calculation, potential of mean force (PMF) calculations were performed as a function of the center-to-center distance between the cancer cell membrane and the gold nanoparticle for all Au+DOX+curcumin systems considered as shown in **Figure 10**. The PMF profiles reveal that the systems containing the neutral Enol and Trans forms of curcumin, namely Au+DOX+Trans and Au+DOX+Enol, which are stable under acidic pH conditions, exhibit substantially stronger binding to cancer cell membrane, with binding free-energies (ΔG) of approximately −30 kcal mol⁻¹. This binding considerably stronger than that of the curcumin-free Au+DOX system, which exhibits a binding free energy of approximately −7 kcal mol⁻¹.

**Figure 10:**
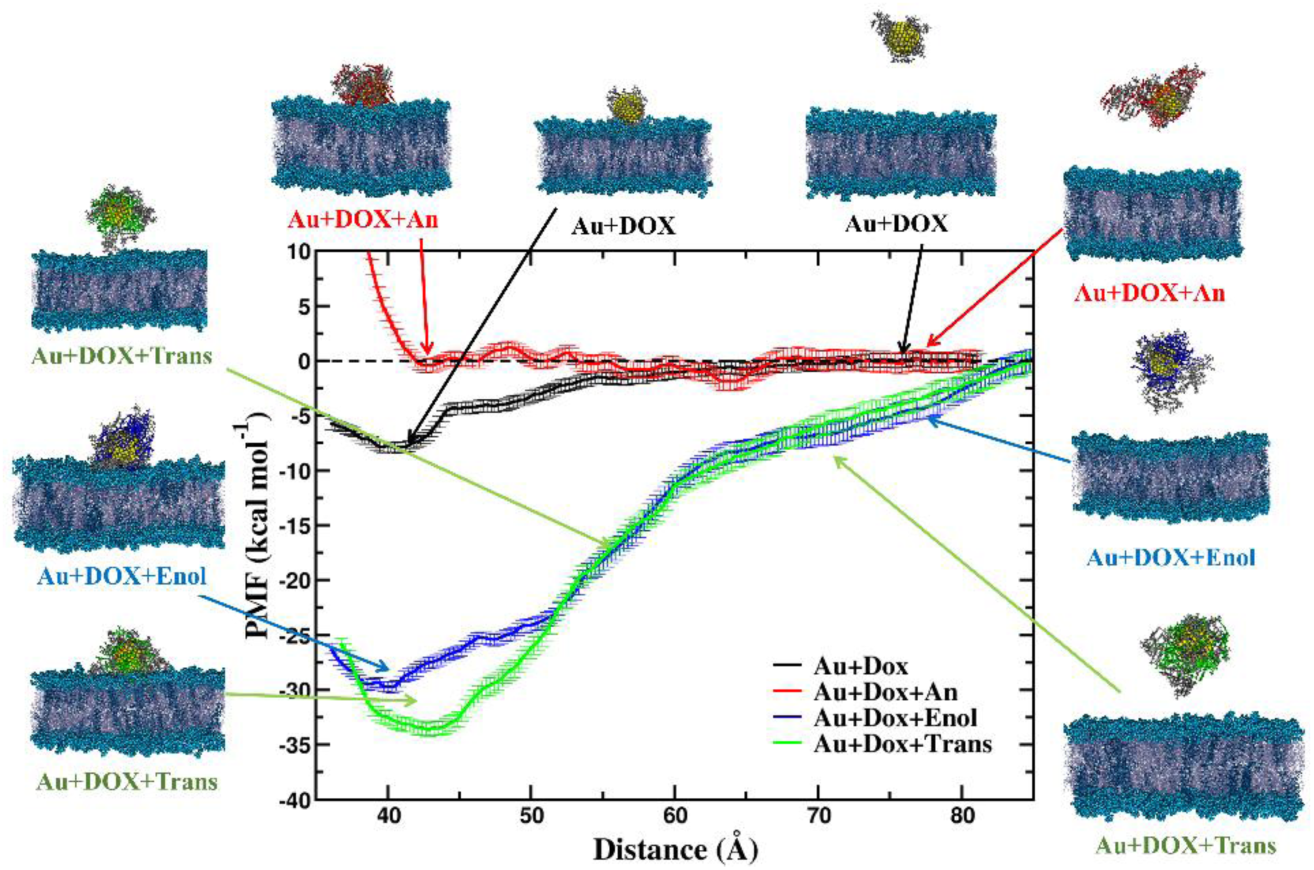
Potential of mean force (PMF) profiles as a function of the center-to-center distance between the cancer cell membrane and the gold nanoparticle were computed for all Au+DOX+curcumin systems, including An, Enol, and Trans forms. The Systems containing neutral curcumin, namely Au+DOX+Trans and Au+DOX+Enol exhibit, exhibit substantially stronger binding to the cancer cell membrane, with a binding energy (ΔG) of approximately −30 kcal mol⁻¹, compared with the curcumin-free Au+DOX system, which shows a binding free energy of approximately −7 kcal mol⁻¹. In contrast, the Au+DOX+An system exhibits very weak binding to the negatively charged cancer membrane.

The enhanced membrane binding of Au+DOX+Trans and Au+DOX+Enol systems can be attributed to the increased number of cationic DOX moieties associated with the gold nanoparticle surface in the presence of curcumin. Consequently, the electrostatic interactions between the positively charged DOX molecules and the negatively charged cancer cell membrane become stronger and extend over a larger separation range. This behaviour consists with the curcumin-free Au+DOX+An system, in which only a limited number of DOX molecules (approximately 18) are associated with the gold nanoparticle surface, resulting in comparatively weaker electrostatic interaction with membrane. In contrast, the presence of negatively charged An curcumin substantially weakens the interaction between the Au+DOX+An complex and the negatively charged cancer cell membrane. The association of An curcumin with the gold nanoparticle surface reduces the effective electrostatic attraction between the complex and the membrane, resulting in very weak binding and nearly negligible binding free energies. These observations are in good agreement with the trends in the MM-PBSA binding energies discussed in the previous section, further supporting the role of curcumin protonation state and DOX surface association in modulating the interaction of the complexes with the cancer cell membrane.

### 14. Association of Niraparib (NIR) within Au+NIR+curcumin complexes and their binding to the cancer cell membrane

To demonstrate that the drug-loading mechanism observed in the Au+DOX+curcumin complexes and their subsequent binding to the cancer cell membrane can be generalized to other cationic anticancer drugs of comparable size, Au+NIR, Au+NIR+An, Au+NIR+Enol, and Au+NIR+Trans complexes were prepared using the same protocol. The corresponding simulation snapshots and structural analyses are presented in **Figure 11**, in order to compare the results obtained for Au+DOX+curcumin complexes.

**Figure 11:**
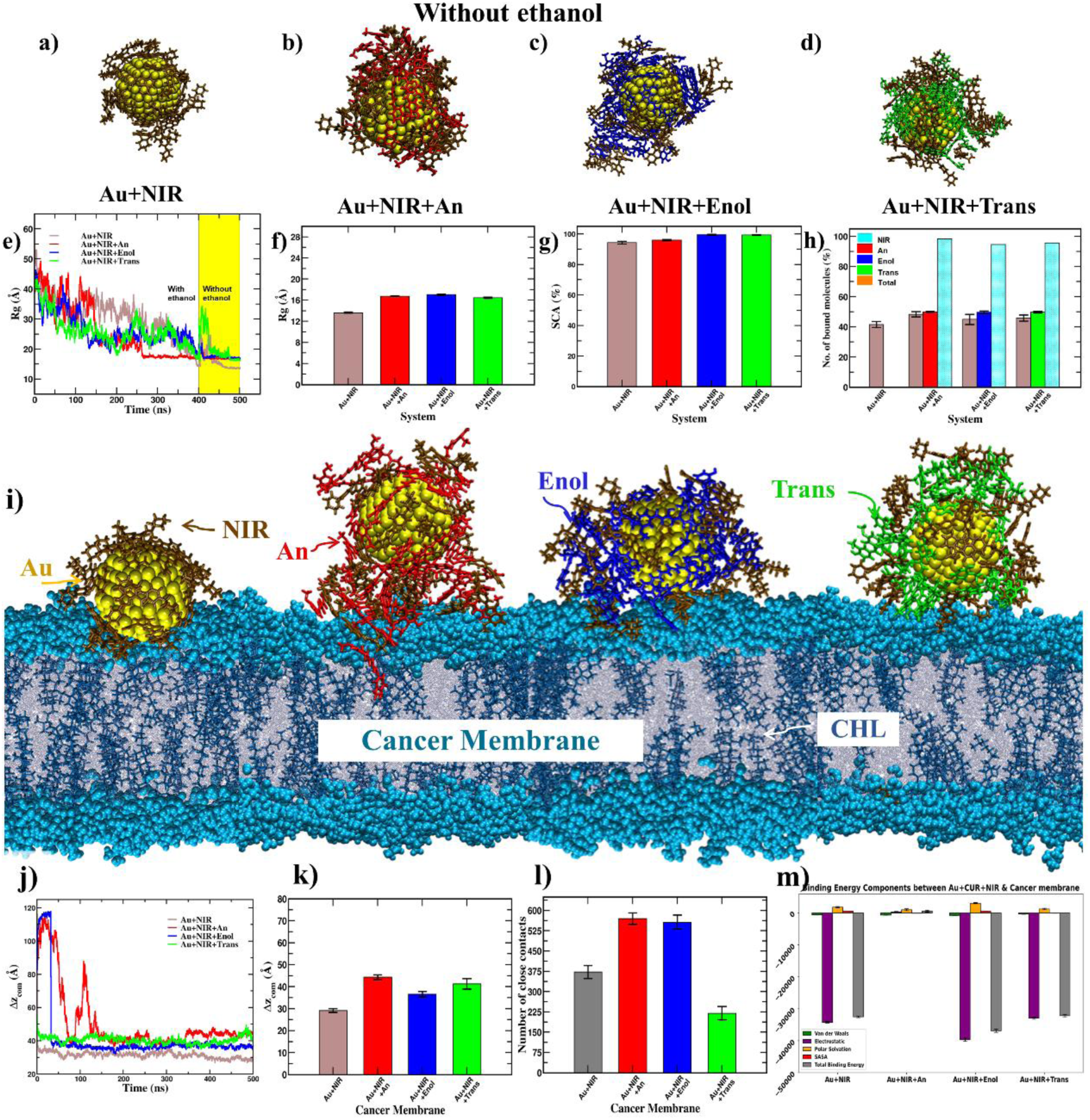
Final snapshots showing curcumin (An, Enol, Trans tautomers) and niraparib (NIR) arrangement on the AuNP surface (ethanol-free) for (a) Au+NIR, (b) Au+NIR+An, (c) Au+NIR+Enol, and (d) Au+NIR+Trans. It can be observed that only 20 of 50 NIR molecules adsorb in the binary Au+NIR system. Whereas, co-loading NIR with curcumin increases NIR adsorption and nanocomposite stability across all curcumin isoforms. This trend is consistent with previously reported doxorubicin–AuNP loading behaviour.

Structural analysis of the Au+NIR+curcumin complexes shows that approximately 20 NIR molecules are adsorbed onto the gold nanoparticle surface, which is comparable to the number of DOX molecules (17–20) adsorbed in the curcumin-free Au+DOX system. Furthermore, analyses of the radius of gyration, surface coverage (SCA), and number of drug molecules associated with the gold nanoparticle demonstrate that NIR loading onto the nanoparticle surface is enhanced in the presence of the curcumin. The Au+NIR+curcumin complexes also adopt a relatively more compact configuration and exhibit greater surface coverage than the corresponding Au+DOX+curcumin complexes.

The binding of the Au+NIR+curcumin complexes to the cancer cell membrane was further examined and compared with that of the corresponding Au+DOX+curcumin complexes. Analyses of the center-to-center distance, number of close-contacts, and MM-PBSA binding energies consistently indicate that the Au+NIR+curcumin complexes exhibit stronger interactions with the cancer cell membrane than the Au+DOX+curcumin complexes. These findings suggest that the curcumin-mediated enhancement of drug loading onto the gold nanoparticle surface is not specific to DOX but can be extended to other cationic anticancer drugs, such as NIR, with comparable molecular dimensions. Moreover, both DOX and NIR exhibit enhanced membrane binding in the presence of the neutral Enol and Trans forms of curcumin. These curcumin forms are favoured under acidic conditions, which are characteristic of the tumour microenvironment. Collectively, these results support the potential generalizability of the proposed Au nanoparticle-curcumin platform for enhanced loading and membrane targeting of cationic anticancer drugs under the tumour-associated acidic conditions.

## Discussion

Ultra-small gold nanoparticles exhibit superior tumour infiltration for enhanced cellular drug uptake, alongside molecular-like optical and electronic properties. Noncovalent adsorption of curcumin preserves its intrinsic anticancer and antioxidant characteristics, rendering the investigation of diverse curcumin forms loaded onto these nanoparticles significantly promising for therapeutic applications and nanoparticle synthesis optimization (*17*). Analysis of Rg, radial density profiles, and curcumin packing demonstrates that neutral Au-curcumin complexes (Au+Enol; Au+Trans), corresponding to acidic to neutral pH conditions, exhibit greater structural compactness with high surface coverage than the deprotonated Au+An from prevalent at alkaline pH. Given the acidic extracellular and alkaline intracellular tumour microenvironment, the enhanced compactness of Au-curcumin complexes under acidic conditions may provide greater shielding against premature drug release, while their reduced compactness under alkaline conditions could facilitate easier curcumin desorption intracellularly (*87*). This pH-responsive behaviour suggests a potential mechanism for sustained and environment-responsive curcumin delivery. Additionally, analysis of curcumin regions reveals that inner-region curcumin molecules form a mechanically robust interfacial shell, while the outer-region curcumin molecules remain flexible with greater solvation. Furthermore, SCA and RMSF profiles together with MMPBSA calculations indicate that the deprotonated curcumin (An) molecules extend outward from the gold nanoparticle surface, reducing SCA to mitigate electrostatic repulsions. Enhanced solvent accessibility in ethanol stabilizes these conformations, limiting curcumin flexibility and increasing intermolecular spacing, which could favour efficient drug loading and releasing dynamics. Transitioning Au-curcumin complexes to more polar solvents (e.g., ethanol to methanol, or pure water) diminishes molecular stacking while strengthening gold-curcumin interactions, yielding smaller, more compact nanostructures (*88*). These compact morphologies could provide structural features to light absorption, electrical conductivity, and self-assembly into ordered conductive frameworks, highlighting their potential relevance to sensors and electronic applications. Analysis of bound water calculations shows that Au-curcumin complexation elevates curcumin hydration levels, which may favour aqueous disposability and potentially contribute to its bioavailability for biomedical applications. This finding consistent with previous experimental studies (*39*). The solvent polarity dependence in connection with hydrogen-bonding capacity is useful in tuning nanoparticle morphology during synthesis. PMF calculations reveal that deprotonated (An) curcumin and its mixture with the Enol form can serve as effective dispersing agents through electrostatic repulsion, potentially promoting colloidal stability, reducing aggregation, and supporting consistent drug-loading and release behaviour for curcumin delivery. Moreover, the inclusion of deprotonated curcumin (An) in these complexes may be important for synthesizing ultra-small gold nanoparticles by leveraging curcumin’s reducing capability (*34,66,68*).

Furthermore, our molecular dynamics (MD) simulations suggest that the loading of doxorubicin (DOX)/niraparib (NIR) onto Au nanoparticles is significantly enhanced in the presence of different curcumin forms. This enhanced loading could potentially reduce the exposure of free DOX/NIR through its association with curcumin-functionalized Au nanoparticles, providing a possible strategy for modulating DOX/NIR-delivery. In addition, these complexes may have potential for improving drug-delivery efficiency and these observations are consistent with recent experimental reports (*89,90*). Under the acidic conditions characteristic of the tumor micro environment, curcumin predominantly adopts neutral forms (Enol and Trans), which facilitate stronger complexation with positively charged DOX/NIR on the Au nanoparticles. In contrast, cancer cell membranes typically possess a net negative charge within the cellular microenvironment, promoting favorable electrostatic interactions with the positively charged Au+DOX/NIR-curcumin complexes. The MD simulation results further reveal that neutral curcumin (Enol and Trans forms) enhances these electrostatic interactions, leading to stronger binding of Au+DOX/NIR+curcumin complexes to the negatively charged model membrane, consistent with conditions associated with an acidic tumor microenvironment. Overall, this study provides molecular-level insights into the structural and intermolecular mechanisms underlying Au+DOX/NIR+curcumin complex formation, DOX/NIR loading, and membrane interactions, highlighting their potential relevance to controlled cancer drug delivery.

## Conclusions

All-atom molecular dynamics simulations were employed over microsecond timescales to elucidate how curcumin tautomerism and solvent environment control the structure and stability of curcumin-coated gold nanoparticles. Followed this, cationic drugs such as doxorubicin (DOX) and niraparib (NIR) were loaded onto the Au-curcumin composites to investigate their loading capacity, and molecular interactions. Finally, the binding of Au+DOX/NIR+curcumin complexes to the cancer cell membrane was demonstrated to investigate differences in their interactions. Their interactions were further compared with a control neutral DMPC membrane. Neutral Enol and Trans-keto curcumin forms generate compact, region-organized coatings with high surface coverage, which may provide enhanced surface protection and favour drug-loading under acidic to near-neutral conditions. In contrasts, the deprotonated Enolate state promotes more extended conformations and reduced surface coverage due to electrostatic repulsion, which could contribute to enhanced colloidal stability and dispersibility under alkaline conditions. Despite this lower surface coverage, binding free energy analysis indicates that enolate-rich complexes exhibit stronger thermodynamic stabilization, highlighting the strong Au-curcumin interactions associated with deprotonation. Region-resolved metrics of size, orientation, and flexibility reveal that inner-region curcumin molecules align parallel to the gold surface and are significantly less flexible than outer-region molecules, establishing a structurally robust interfacial shell. Replacing ethanol with methanol reduces π–π stacking, enhances direct Au-curcumin contacts, and increases surface coverage for all curcumin forms, resulting in smaller and more compact nanostructures. Potential of mean force calculations demonstrate that enolate curcumin can nearly completely screen Au–Au attraction and extend the effective interaction range, while neutral and mixed systems provide partial screening. These findings provide molecular-level insight into how deprotonated curcumin may contribute both to nanoparticle formation and to colloidal stabilization in solution.

The MD simulations of Au+DOX/NIR and Au+DOX/NIR+curcumin complexes reveal that DOX/NIR molecules associate with curcumin through stacking interactions, enhancing their loading onto the Au nanoparticles by reducing electrostatic repulsion. This well-organized DOX/NIR-curcumin complexation may provide a favourable structural basis for drug incorporation within the nanoparticle assembly. Furthermore, curcumin molecules preferentially occupy the first coordination layer on the Au nanoparticle surface due to stronger van der Waals interactions compared to those between DOX/NIR with Au atoms. In the Au+DOX/NIR+An complex, DOX/NIR molecules are dispersed across the surface and screened by negative charged An molecules, and adopting more extended conformations. However, this system exhibits weaker interactions with the negatively charged model membrane. In contrast, DOX/NIR loaded Au nanoparticle in presence of neutral curcumin (Enol and Trans forms) form compact structures with a net positive surface charge, leading to stronger interactions with the negatively charged head groups of the model membranes. PMF calculations, together with the relative changes in MM-PBSA binding energies, further indicate that Au+DOX+Enol and Au+DOX+Trans complexes exhibit enhanced membrane binding compared to Au+DOX when interacting with the negatively charged model cancer membrane. These electrostatically driven binding interactions were further compared with those observed for the control neutral DMPC membrane, providing a basis for distinguishing charge-dependent membrane association.

Overall, this work provides a molecular-level framework for selecting curcumin protonation states and solvent conditions to precise control morphology, dispersion, and interfacial hydration in curcumin–gold nanocomposites. It also provides molecular insights into the enhanced loading and membrane interactions of cationic anticancer drugs such as doxorubicin and niraparib in the presence of curcumin on Au nanoparticles. These findings may contribute to the rational design of curcumin-based nanocarriers and functional gold nanoclusters for applications in catalytic, sensing, and pH-responsive drug delivery systems.

## Supporting information

Curcumin_Gold_Nanocomposites_supportinfo_V2

## AUTHOR INFORMATION

### Author contributions

A.G. conceived the work, performed MD, and analysed data. H.N. and SG.N. modelled Au nanoparticles and assisted with analysis. S.B. performed MD for nanocomposites-membrane interactions and related analyses. J.K.K. and S.K. conceptualized, and designed the work. A.G., J.K.K. and S.K. contributed to discussions, data compilation, manuscript writing and revision. All authors approved the final manuscript.

## Acknowledgements

We thank the SAI-HPC, COSMOS Lab (CRIF) and DMACS at Sri Sathya Sai Institute of Higher Learning for providing the necessary computational resources. The authors also acknowledge the assistance of AI large-language-model tools in improving the grammar, clarity, and readability of the manuscript. All scientific content, analysis, and conclusions are solely the responsibility of the authors. The authors are grateful to Bhagawan Sri Sathya Sai Baba, Founder Chancellor, SSSIHL for his constant inspiration.

## Table of Contents (TOC)

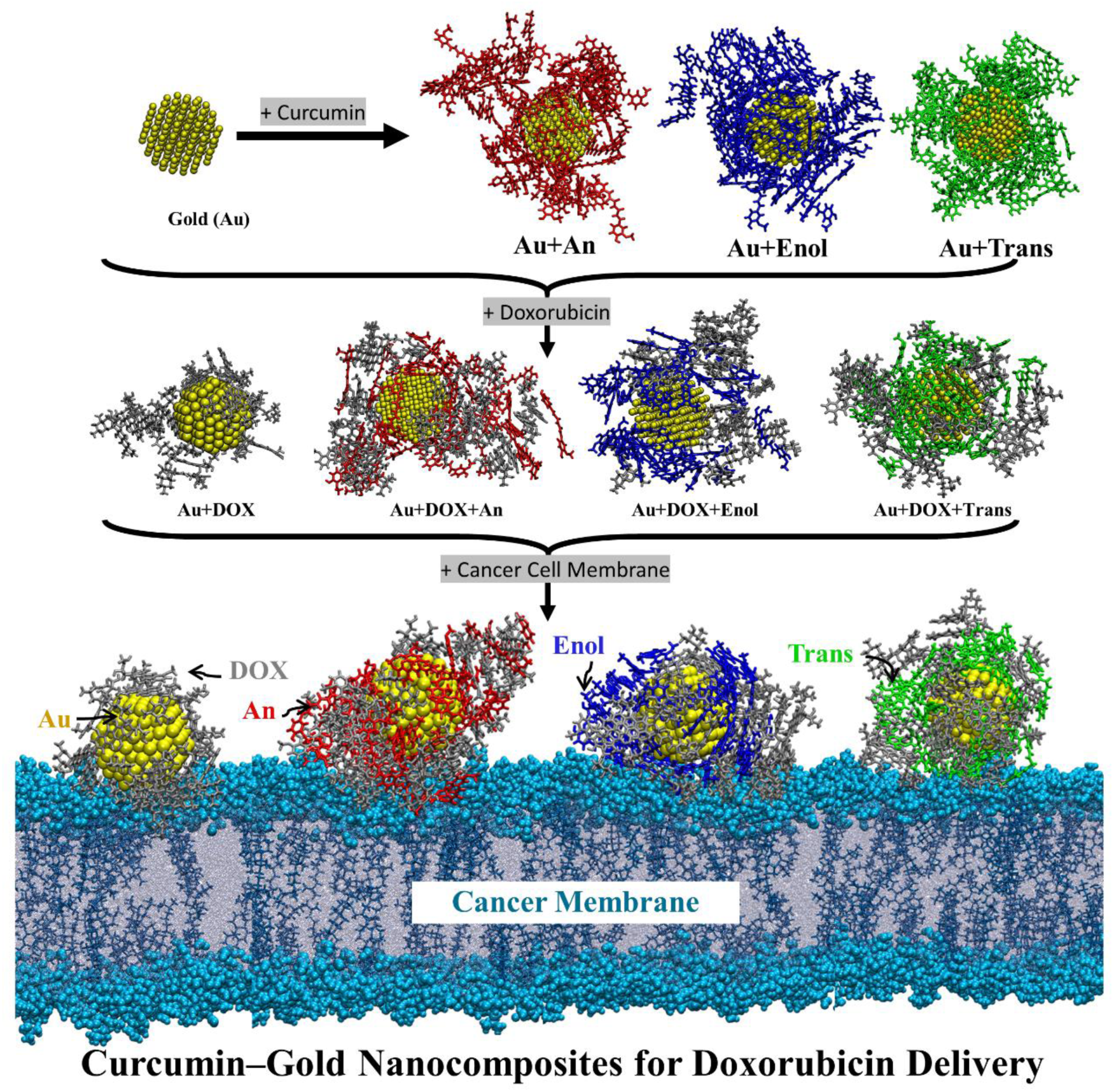

