## Supplementary material for "Curcumin–Gold Nanocomposites for Enhanced Cationic Anticancer Drug Delivery: Molecular Mechanisms of Loading and Membrane Interactions": Curcumin_Gold_Nanocomposites_supportinfo_V2

* Corresponding author

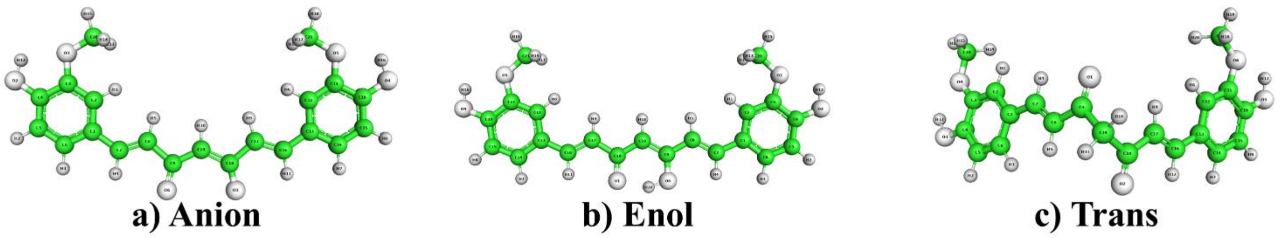

**Figure S1:** Structures of curcumin states: (a) Anion, (b) Enol, and (c) Trans. Partial atomic charges were calculated using Gaussian and AmberTools.

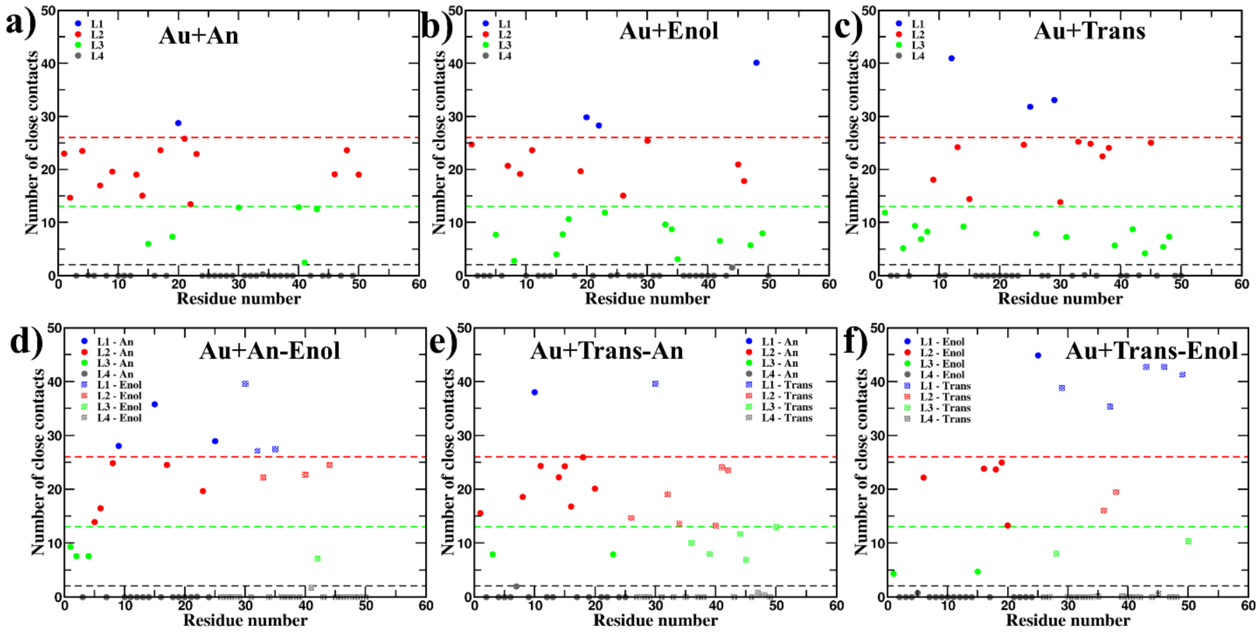

**Figure S2.** Number of close contacts (curcumin atoms within 4 Å of the gold nanoparticle) calculated for: (a) Au+An, (b) Au+Enol, (c) Au+Trans, and mixed curcumin systems (d) Au+An-Enol, (e) Au+Trans-An, and (f) Au+Trans-Enol. For each system, the number of close contacts is determined for every curcumin molecule. This analysis distinguishes molecules that are closest to the gold nanoparticle, partially connected, or not connected at all. Based on these contact counts, curcumin molecules are assigned to different regions (L1, L2, L3 and L4), reflecting their interaction with the gold nanoparticle.

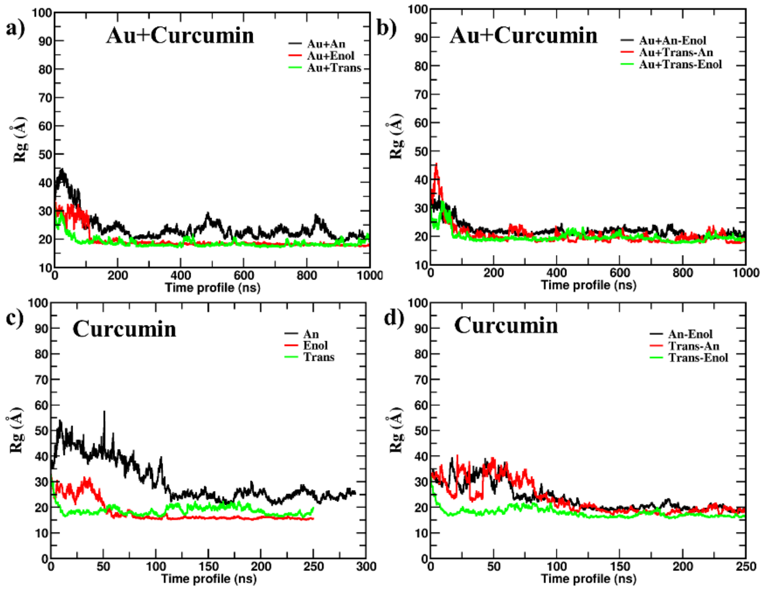

**Figure S3:** Radius of gyration ($R_g$) profiles for: (a) pure curcumin with gold nanoparticle (Au+An, Au+Enol, Au+Trans), (b) mixture curcumin with gold nanoparticle (Au+An-Enol, Au+Trans-An, Au+Trans-Enol), (c) pure curcumin only (An, Enol, Trans), and (d) mixture curcumin only (An-Enol, Trans-An, Trans-Enol). The charged curcumin systems (An and Au+An) require a longer time to reach equilibration compared to the neutral systems (Trans and Enol), as reflected by the delayed stabilization of their Rg profiles.

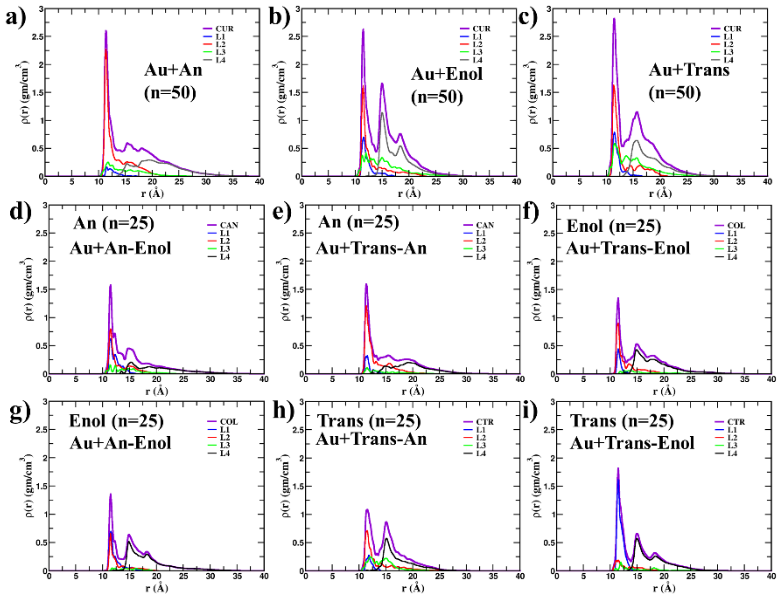

**Figure S4:** Density profiles of curcumin molecules around the gold nanoparticle for pure systems (a) Au+An, (b) Au+Enol, (c) Au+Trans and mixture systems (d & g) Au+An-Enol, (e & h) Au+Trans-An, (f & i) Au+Trans-Enol. The first peak in each density profile arises predominantly from molecules in regions L1 and L2 molecules, which are closest to the gold nanoparticle. In contrast, the second peak and the extended tail region derive from molecules in the outer regions, L3 and L4 molecules. In mixture systems containing An, the anion (An) curcumin contributes exclusively to the first peak, while the neutral counterparts (Enol, Trans) contributes to the first peak and are responsible for the second peak. This indicates that An curcumin retains its distinct interaction pattern even within mixtures.

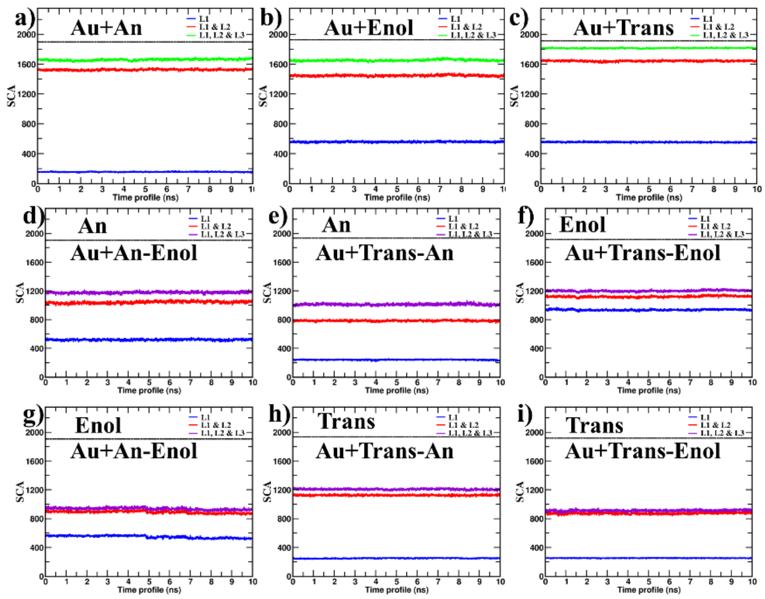

**Figure S5:** Surface coverage area (SCA) of the gold nanoparticle in the presence of pure curcumin systems—(a) Au+An, (b) Au+Enol, (c) Au+Trans and mixture systems (d & g) Au+An-Enol, (e & h) Au+Trans-An, (f & i) Au+Trans-Enol. Among all systems, Au+Trans exhibits the highest surface coverage from combined regions (L1, L2, and L3) molecules. The surface coverage contributed by each region is directly proportional to the number of curcumin molecules occupying that region. Notably, the contribution from L3 to the overall SCA is minimal, since only a small number of atoms from L3 region molecules are in direct contact with the gold nanoparticle.

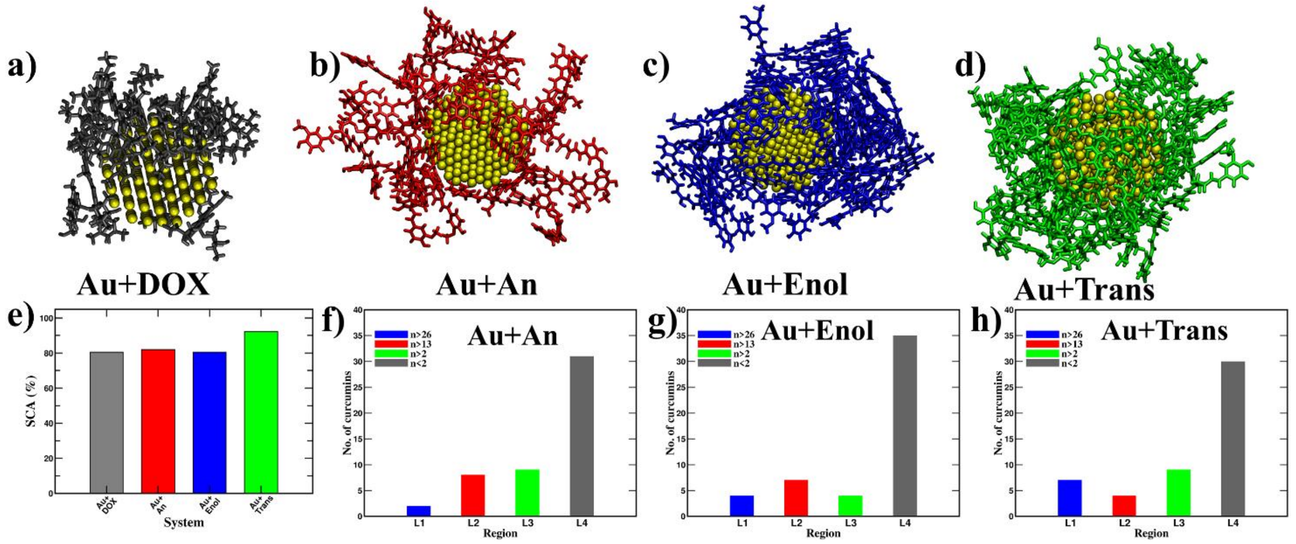

**Figure S6:** Final simulation snapshots of Au+DOX and Au+Curcumin (An, Enol and Trans) in complex with the gold nanoparticle in the presence of $Na^+$ counter ions. Consistent with the trend observed under $H^+$, DOX and the An curcumin show a lower tendency to aggregate around the gold nanoparticle, whereas the trans and enol forms readily wrap around the nanoparticle surface. In all the complexes, curcumin coverage around Au nanoparticle is comparatively **lower** in the presence of $Na^+$ than $H^+$. Whereas, DOX coverage remains similar between the two ionic conditions, indicating an overall consistent trend across ion types.

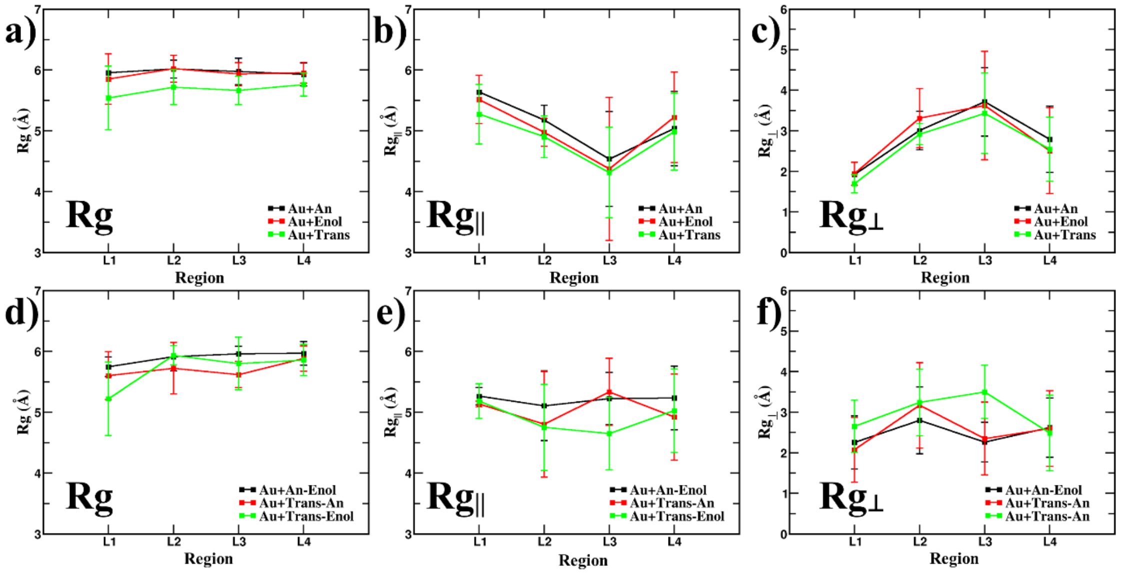

**Figure S7:** Radius of gyration (Rg) calculated for each region of curcumin molecules in gold nanoparticle systems: (a and d) total Rg; (b and e) parallel component of Rg ($Rg_{||}$); (c and f) normal component of Rg ($Rg_{\perp}$). For pure systems, Au+An and Au+Enol show similar values for Rg, while trans Au+Trans exhibits a lower Rg compared to other two. Rg remains nearly constant across regions, indicating that the compactness of curcumin molecules is independent of their position relative to the nanoparticle. In contrast, $Rg_{||}$ decreases from L1 to L3, suggesting a reduction in molecule extension parallel to the surface, while $Rg_{\perp}$ increases over the same range. In mixture systems, deviations in Rg, $Rg_{||}$, and $Rg_{\perp}$ from L1 to L4 are smallest for Au+An-Enol, suggesting the most homogeneous region-dependent structure in this composition.

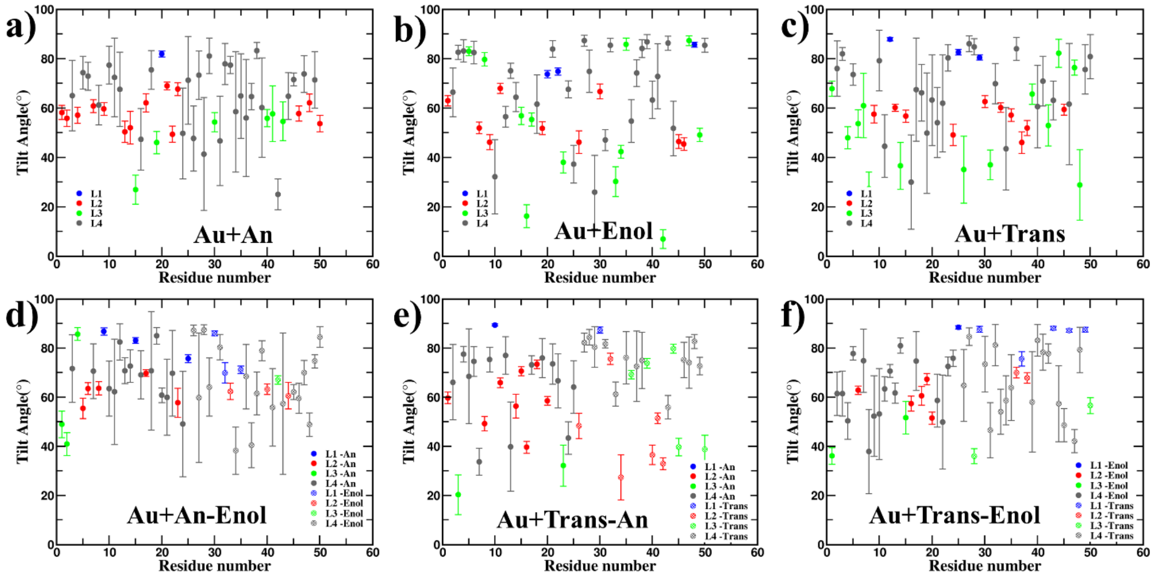

**Figure S8:** Tilt angles of curcumin molecules with respect to the surface normal of the gold nanoparticle are shown for (a) Au+An, (b) Au+Enol, (c) Au+Trans (d) Au+An-Enol, (e) Au+Trans-An, (f) Au+Trans-Enol cases. In all systems, the average tilt angle decreases from L1 to L3, indicating that L1 molecules lie most parallel to the surface. As L4 molecules are not in direct contact with the gold surface, their tilt angles show greater variability and can adopt any orientation.

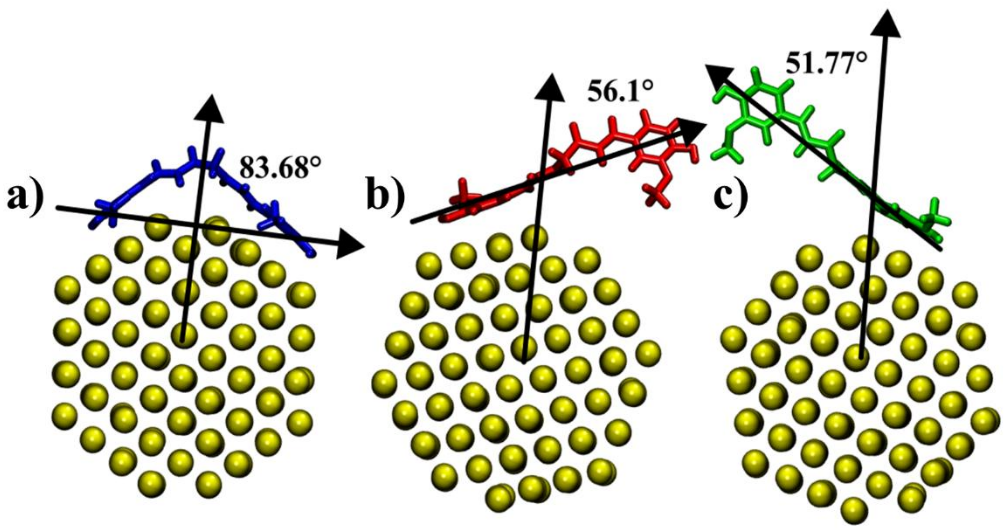

**Figure S9:** Visualization of curcumin molecules in each region: (a) L1, (b) L2, and (c) L3. In L1, all atoms of the curcumin molecules are connected to the gold nanoparticle, indicating complete surface contact. In L2, approximately half of the atoms in each curcumin molecule are directly connected to the gold surface, reflecting partial contact. In L3, only a few atoms of each curcumin molecule are associated with the nanoparticle, signifying minimal direct interaction.

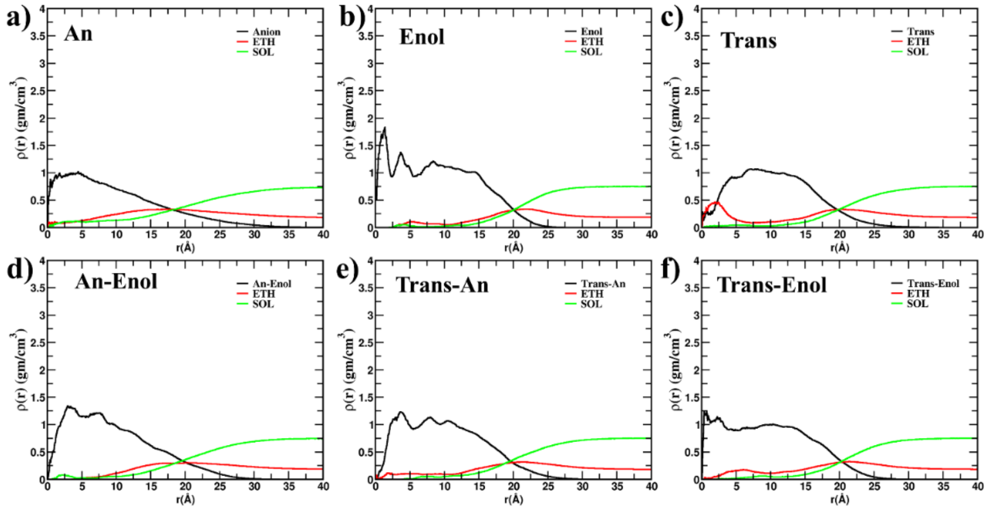

**Figure S10:** Density profiles of curcumin molecules in the absence of a gold nanoparticle for pure systems—(a) An, (b) Enol, (c) Trans and mixture systems (d) An-Enol, (e) Trans-An, (f) Trans-Enol. The profiles reveal that the An (anion) and An-Enol mixture exhibit noticeably longer tail regions compared to other cases, indicating that these systems have a more extended molecular distribution in the absence of the gold nanoparticle.

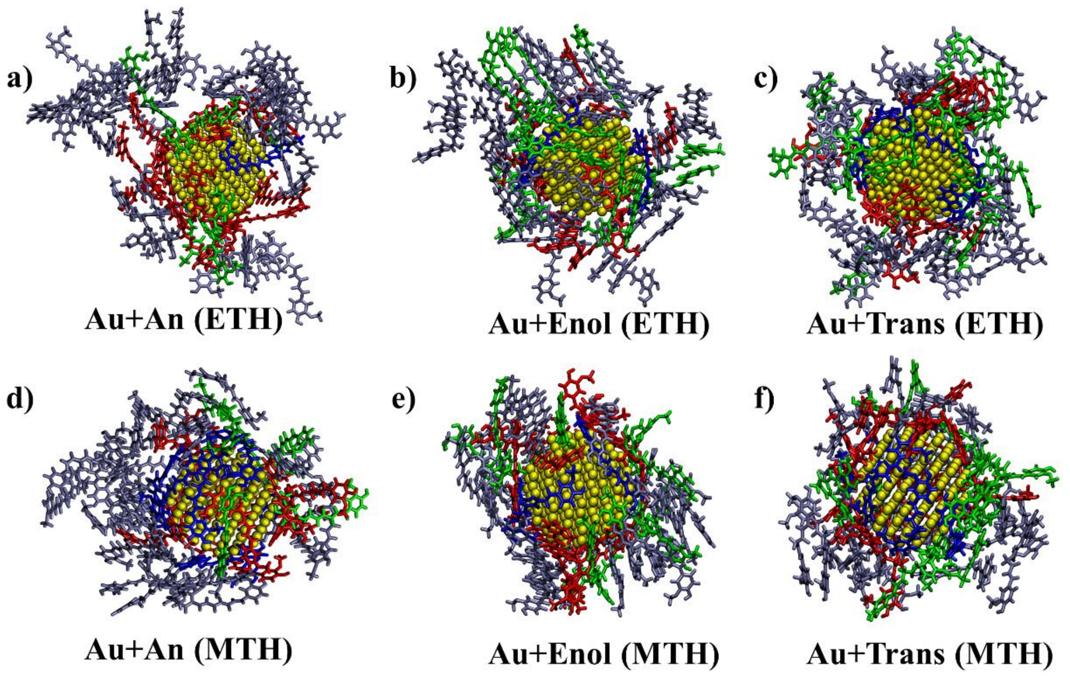

**Figure S11:** Final simulation snapshots of pure curcumin molecules-(a) An, (b) Enol, and (c) Trans in the presence of ethanol and (d) An, (e) Enol, and (f) Trans in the presence of methanol, each wrapped around a gold nanoparticle. Curcumin molecules are color-coded by the number of contacts with the nanoparticle: blue corresponds to the highest number of contacts, followed by red and green, while grey indicates molecules with no contact.

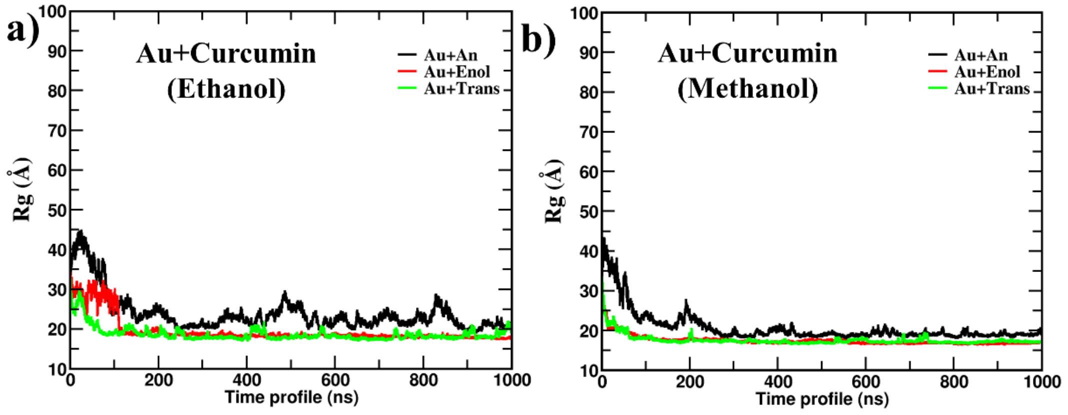

**Figure S12:** Radius of gyration (Rg) profiles for curcumin molecules with a gold nanoparticle in the presence of (a) ethanol and (b) methanol. The charged curcumin system (Au+An) requires a longer equilibration time than the neutral systems (Au+Trans and Au+Enol), as indicated by the delayed stabilization of its Rg profile. Additionally, the trans (Au+Trans) and enol (Au+Enol) systems exhibit smaller Rg values than the anion (Au+An), reflecting their more compact molecular sizes.

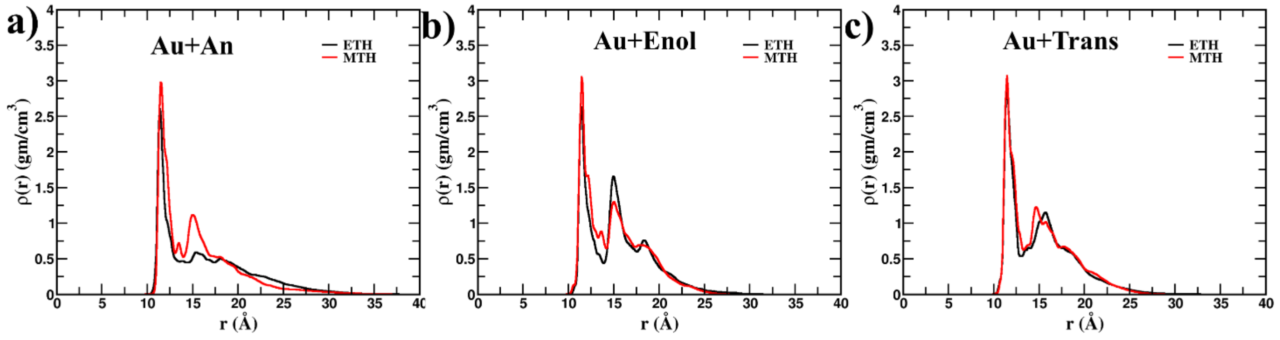

**Figure S13:** Density profiles for pure curcumin species—(a) An, (b) Enol, and (c) Trans around the gold nanoparticle in the presence of ethanol and methanol. In all systems, the methanol environment produces a higher and sharper density peak compared to ethanol, indicating tighter molecular packing of curcumin molecules around the nanoparticle in methanol than in ethanol.

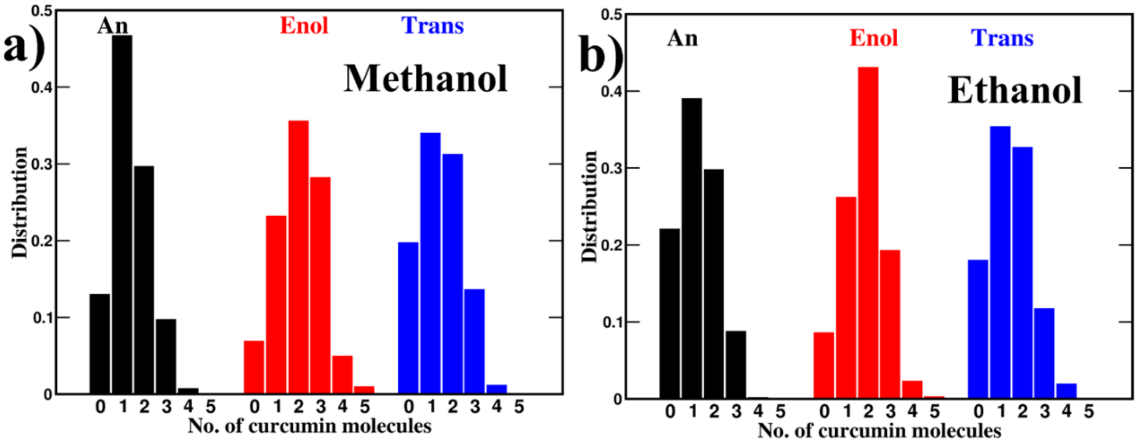

**Figure S14:** Probability distribution comparison of curcumin molecules around the gold nanoparticle in the presence of different solvents-ethanol and methanol. A significant difference in the distribution is observed for the anion system (Au+An), whereas the trans system (Au+Trans) shows minimal change between the two solvents.

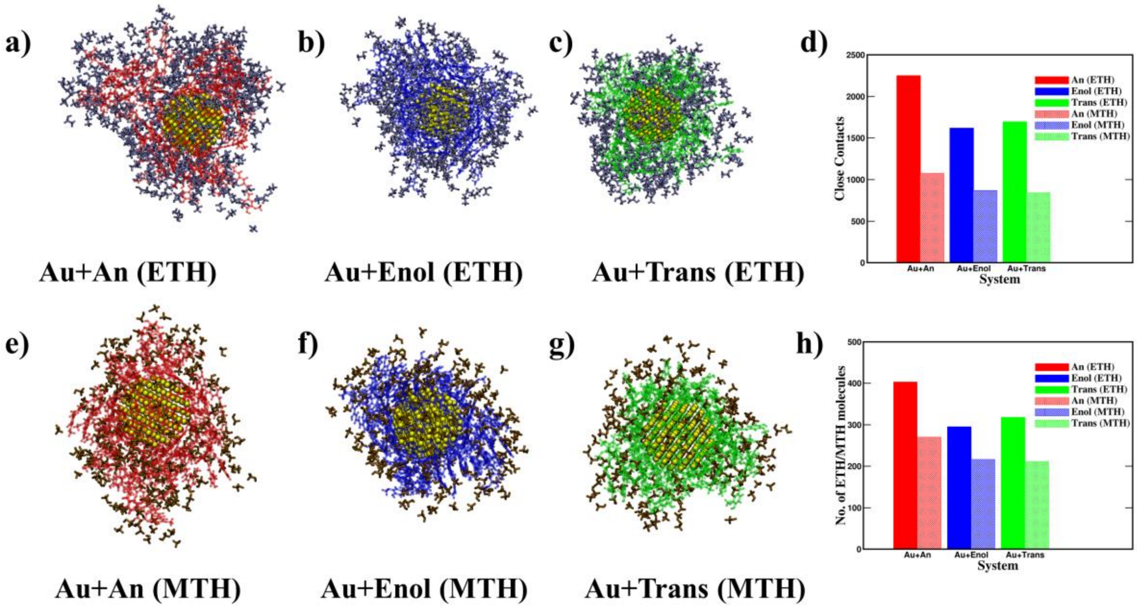

**Figure S15:** The ethanol (a–c) and methanol (e–g) molecules surrounding curcumin in the Au-curcumin complexes were visualized for the respective ethanol and methanol solvated systems. The number of close contacts between curcumin and the ethanol/methanol solvent molecules (d), as well as the number of ethanol/methanol molecules located within 5 Å of curcumin (h) were compared. A higher number of ethanol molecules were observed in the vicinity of the Au–curcumin complex compared with the corresponding methanol-solvated complexes. This enhanced ethanol solvation may be attributed to the relatively greater hydrophobicity of ethanol compared with methanol, which promotes closer association with the hydrophobic moieties of curcumin and thereby contributes to improved solvation and stabilization of the complex.

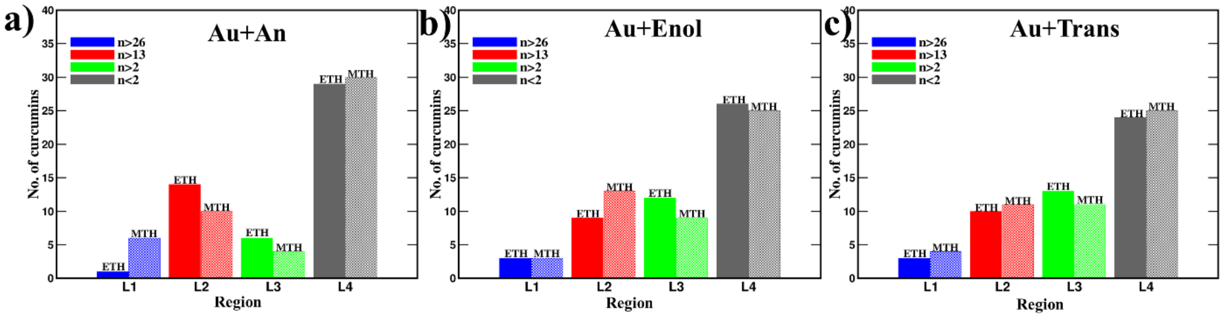

**Figure S16:** Histograms displaying the number of contacts per curcumin molecule for (a) An, (b) Enol, and (c) Trans, compared in environments containing ethanol and methanol. The distributions reveal that in the outer regions (L3 and L4), curcumin molecules behave similarly regardless of the solvent. However, In the inner regions (L1 and L2) near the gold nanoparticle, more curcumin molecules are present in methanol than in ethanol, indicating stronger gold–curcumin binding in methanol.

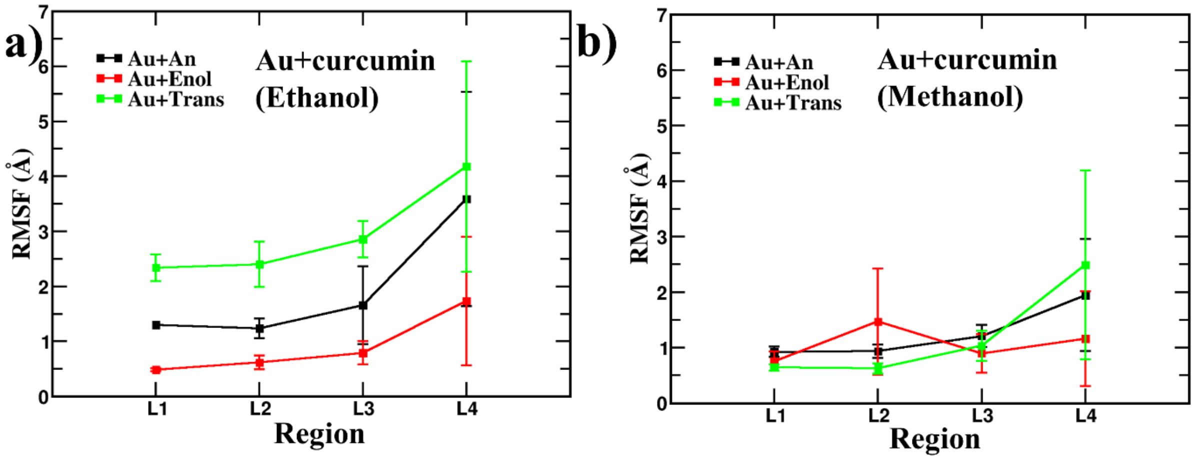

**Figure S17:** Root mean square fluctuation (RMSF) profiles for gold–curcumin complexes in the presence of (a) ethanol and (b) methanol. The Au+An system exhibits higher flexibility compared to the Au+Trans system in the innermost regions. Additionally, the Au+Enol shows increased rigidity in regions L3 and L4, indicating more tightly packed molecular arrangements in these outer regions.

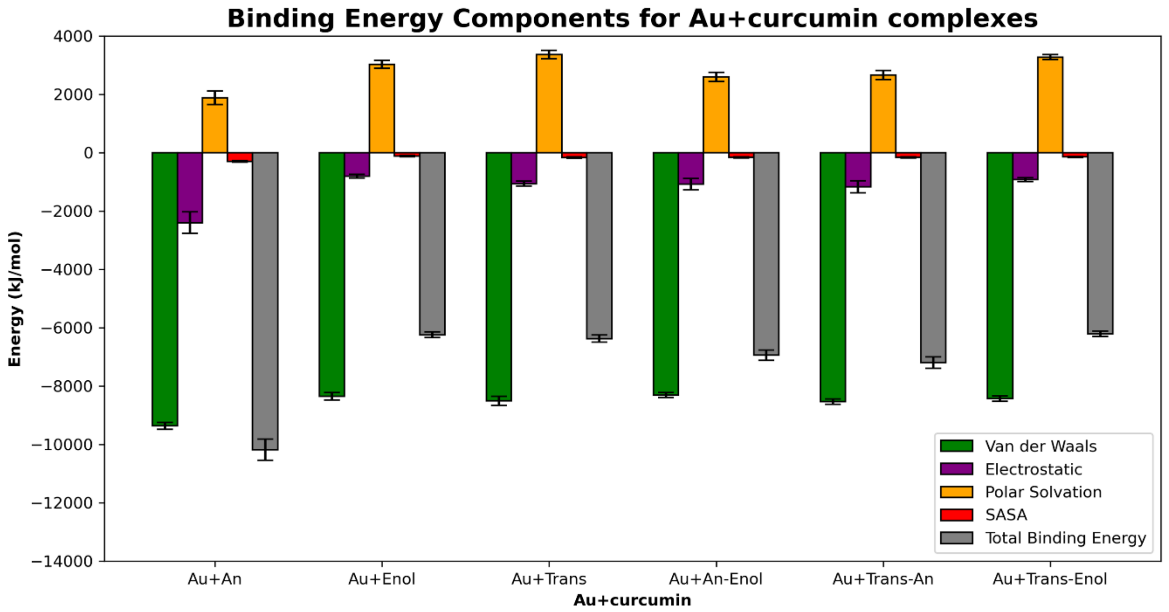

**Figure S18:** Individual contributions to the total binding free energy arising from Au-curcumin, Au-ethanol, and curcumin-ethanol interactions, are analysed to elucidate binding differences in Au-curcumin nanocomposites.

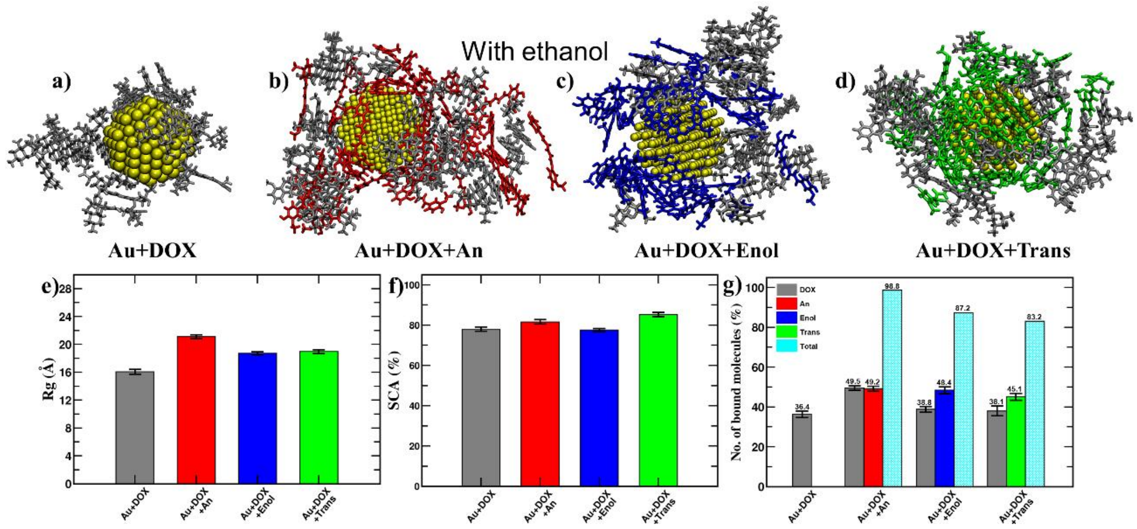

**Figure S19:** Final snapshots illustrating doxorubicin (DOX) loading onto the Au nanoparticle surface: (a) Au+DOX in the absence of curcumin, and (b) Au+DOX+An, (c) Au+DOX+Enol, and (d) Au+DOX+Trans in the presence of curcumin under ethanol solvent conditions.

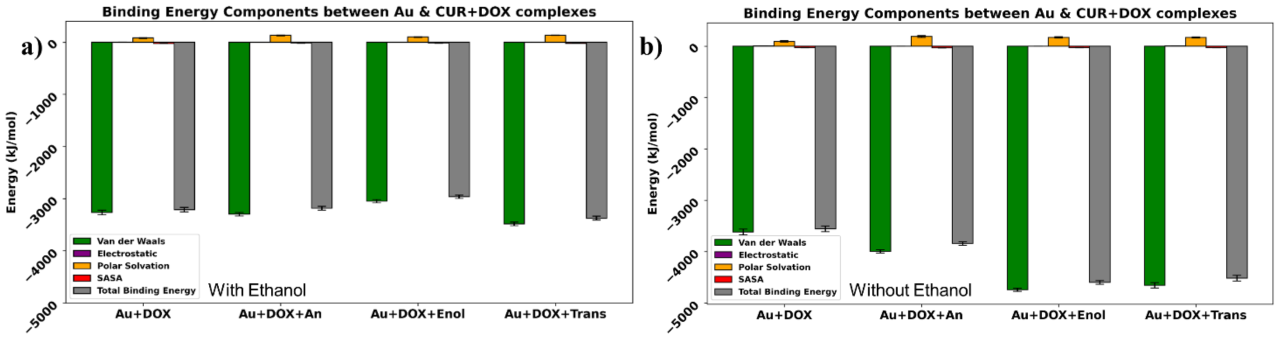

**Figure S20:** Difference in binding energy contributions between Au and DOX+curcumin components across various Au+DOX+curcumin complexes under (a) ethanol and (b) ethanol-free solvent conditions.

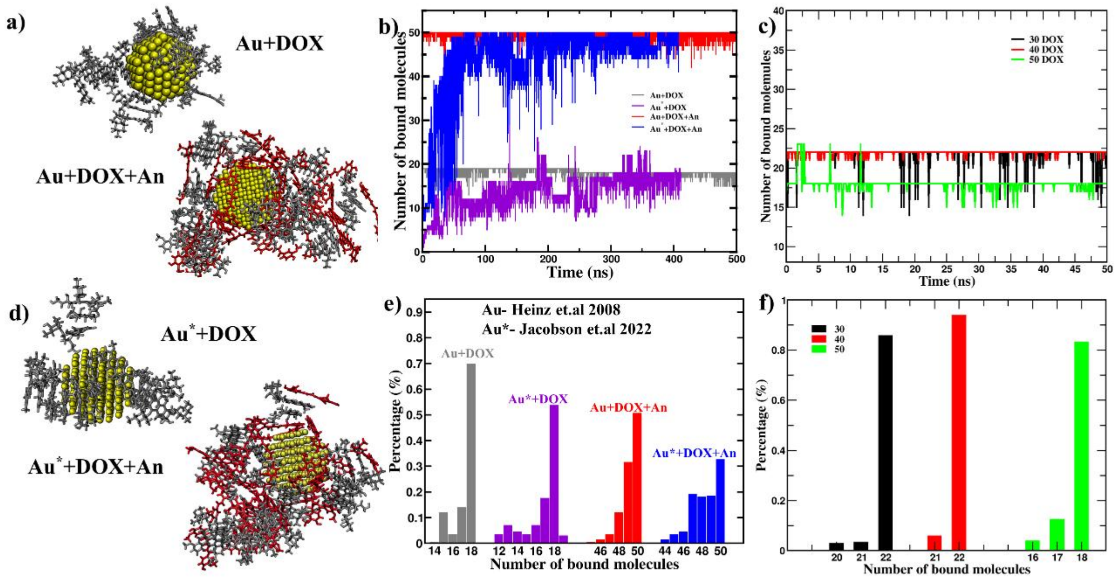

**Figure S21:** Simulation snapshots of DOX-loaded Au and Au+An-curcumin complexes using (a) the original force field employed in this study (Au, Heinz et al 2008) and (d) the alternative gold force field reported by Jacobson et al (Au* 2022) for compassion. (b) Time evolution and (e) corresponding distributions of the number of DOX molecules loaded onto Au surface. (c) Time evolution and (f) corresponding distributions of the number of DOX molecules loaded onto the Au surface for different initial number of DOX molecules.

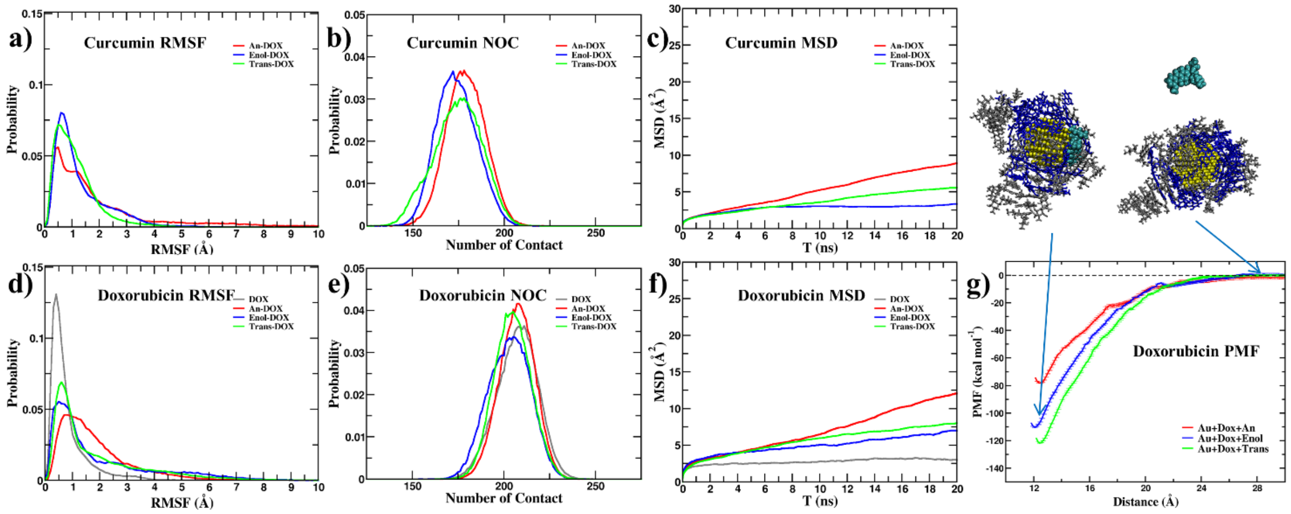

**Figure S22:** The RMSF distributions (a, d), number of intermolecular contacts (NOC) (b, e), and mean-square-displacement MSD (c, f) of curcumin and doxorubicin (DOX) were analysed to assess molecular stability within the Au+DOX+curcumin complexes. Comparison of (g) PMF profiles between DOX and Au of the complexes as a function of center-to-center distance. Au+DOX+An shows the highest conformational flexibility and translational mobility, whereas Au+DOX+Enol is the most conformational restrained, indicating greater complex stability. In contrast, curcumin-free Au+DOX exhibits markedly lower RMSF and MSD values, suggesting that curcumin enhances DOX mobility irrespective of tautomeric form. The higher NOC and MSD for Au+DOX+An further indicate a dynamic, exchange-prone interface.

| TRANS | | | ENOL | | | AN | | |
|---|---|---|---|---|---|---|---|---|
| C1 | ca | 0.061543 | C1 | ca | 0.036613 | C1 | ca | 0.099423 |
| C2 | ca | -0.236025 | C2 | ca | -0.190315 | C2 | ca | -0.21136 |
| C3 | ca | 0.181968 | C3 | ca | 0.140684 | C3 | ca | 0.144716 |
| C4 | ca | 0.260278 | C4 | ca | 0.279416 | C4 | ca | 0.240799 |
| C5 | ca | -0.224395 | C5 | ca | -0.207971 | C5 | ca | -0.22597 |
| C6 | ca | -0.216226 | C6 | ca | -0.242958 | C6 | ca | -0.278602 |
| H1 | ha | 0.145154 | H1 | ha | 0.142138 | H1 | ha | 0.137085 |
| H2 | ha | 0.183809 | H2 | ha | 0.179769 | H2 | ha | 0.168825 |
| H3 | ha | 0.165075 | H3 | ha | 0.161661 | H3 | ha | 0.162509 |
| C7 | ce | -0.016144 | C7 | ce | -0.011702 | C7 | ce | -0.061103 |
| H4 | ha | 0.118922 | H4 | ha | 0.144177 | H4 | ha | 0.103971 |
| C8 | cf | -0.38948 | C8 | cf | -0.408971 | C8 | cf | -0.4659 |
| H5 | ha | 0.17191 | H5 | ha | 0.174068 | H5 | ha | 0.178602 |
| C9 | c | 0.63265 | C9 | cf | 0.67681 | C9 | c | 0.863779 |
| O1 | o | -0.545347 | C10 | ca | 0.256998 | C10 | ca | 0.240799 |
| C10 | ca | 0.260278 | C11 | ca | 0.162099 | C11 | ca | 0.144716 |
| C11 | ca | 0.181968 | C12 | ca | -0.208403 | C12 | ca | -0.21136 |
| C12 | ca | -0.236025 | C13 | ca | 0.063784 | C13 | ca | 0.099423 |
| C13 | ca | 0.061543 | C14 | ca | -0.253347 | C14 | ca | -0.278602 |
| C14 | ca | -0.216226 | C15 | ca | -0.200835 | C15 | ca | -0.22597 |
| C15 | ca | -0.224395 | H6 | ha | 0.142389 | H6 | ha | 0.137085 |
| H6 | ha | 0.145154 | H7 | ha | 0.166168 | H7 | ha | 0.162509 |
| H7 | ha | 0.165075 | H8 | ha | 0.179117 | H8 | ha | 0.168825 |
| H8 | ha | 0.183809 | C16 | ce | -0.009047 | C16 | ce | -0.061103 |
| C16 | ce | -0.016144 | C17 | cf | -0.450626 | C17 | cf | -0.4659 |
| C17 | cf | -0.38948 | H9 | ha | 0.177643 | H9 | ha | 0.178602 |
| H9 | ha | 0.17191 | C18 | c | 0.787477 | C18 | cf | 0.863779 |
| C18 | c | 0.63265 | C19 | ce | -0.801963 | C19 | ce | -0.946457 |
| C19 | c3 | -0.364822 | H10 | ha | 0.212733 | H10 | ha | 0.146871 |
| H10 | hc | 0.100858 | H11 | ha | 0.132411 | H11 | ha | 0.103971 |
| H11 | hc | 0.100858 | O1 | o | -0.645701 | O1 | o | -0.663252 |
| H12 | ha | 0.118922 | O2 | oh | -0.530615 | O2 | oh | -0.558837 |
| O2 | o | -0.545347 | H12 | ho | 0.378633 | H12 | ho | 0.366563 |
| O3 | oh | -0.518433 | O3 | os | -0.241592 | O3 | os | -0.254921 |
| H13 | ho | 0.376649 | C20 | c3 | -0.131665 | C20 | c3 | -0.118765 |
| O4 | os | -0.249633 | H13 | h1 | 0.101473 | H13 | h1 | 0.090744 |
| C20 | c3 | -0.141554 | H14 | h1 | 0.101473 | H14 | h1 | 0.090744 |
| H14 | h1 | 0.106944 | H15 | h1 | 0.101473 | H15 | h1 | 0.090744 |
| H15 | h1 | 0.106944 | O4 | oh | -0.525599 | O4 | oh | -0.558837 |
| H16 | h1 | 0.106944 | H16 | ho | 0.378788 | H16 | ho | 0.366563 |
| O5 | oh | -0.518433 | O5 | os | -0.244744 | O5 | os | -0.254921 |
| H17 | ho | 0.376649 | C21 | c3 | -0.134076 | C21 | c3 | -0.118765 |
| O6 | os | -0.249633 | H17 | h1 | 0.101922 | H17 | h1 | 0.090744 |
| C21 | c3 | -0.141554 | H18 | h1 | 0.101922 | H18 | h1 | 0.090744 |
| H18 | h1 | 0.106944 | H19 | h1 | 0.101922 | H19 | h1 | 0.090744 |
| H19 | h1 | 0.106944 | O6 | oh | -0.624045 | O6 | o | -0.663252 |
| H20 | h1 | 0.106944 | H20 | ho | 0.480413 | | | |

**Table S1:** Charge info of anion, enol and trans curcumins

| Doxorubicin | | | Niraparib | | |
|---|---|---|---|---|---|
| **O1** | os | -0.359 | O1 | o | -0.599 |
| O2 | os | -0.428 | N1 | n4 | -0.192 |
| O3 | oh | -0.738 | N2 | na | 0.293 |
| O4 | oh | -0.631 | N3 | nc | -0.561 |
| O5 | oh | -0.488 | N4 | n | -0.973 |
| O6 | oh | -0.509 | C1 | c3 | 0.062 |
| O7 | o | -0.434 | C2 | c3 | -0.057 |
| O8 | oh | -0.667 | C3 | c3 | -0.174 |
| O9 | o | -0.478 | C4 | c3 | -0.041 |
| O10 | o | -0.466 | C5 | c3 | -0.065 |
| O11 | os | -0.361 | C6 | ca | 0.049 |
| N1 | n4 | -0.546 | C7 | ca | -0.209 |
| C1 | c3 | 0.215 | C8 | ca | -0.209 |
| C2 | c3 | 0.432 | C9 | ca | 0.179 |
| C3 | c3 | -0.079 | C10 | ca | -0.162 |
| C4 | c3 | 0.005 | C11 | ca | -0.162 |
| C5 | ca | -0.196 | C12 | ca | 0.198 |
| C6 | ca | -0.171 | C13 | cc | -0.404 |
| C7 | c3 | 0.343 | C14 | ca | 0.393 |
| C8 | c3 | -0.398 | C15 | ca | -0.302 |
| C9 | c3 | 0.338 | C16 | ca | -0.288 |
| C10 | c3 | -0.017 | C17 | ca | -0.040 |
| C11 | c3 | 0.388 | C18 | ca | -0.146 |
| C12 | ca | 0.367 | C19 | c | 0.844 |
| C13 | c | 0.172 | H1 | hc | 0.044 |
| C14 | ca | 0.329 | H2 | hc | 0.068 |
| C15 | ca | -0.284 | H3 | hc | 0.068 |
| C16 | ca | -0.191 | H4 | hx | 0.137 |
| C17 | c3 | -0.492 | H5 | hx | 0.137 |
| C18 | c3 | 0.427 | H6 | hc | 0.065 |
| C19 | c | 0.613 | H7 | hc | 0.065 |
| C20 | c | 0.647 | H8 | hx | 0.115 |
| C21 | ca | -0.054 | H9 | hx | 0.115 |
| C22 | ca | -0.244 | H10 | hn | 0.300 |
| C23 | ca | 0.278 | H11 | ha | 0.149 |
| C24 | ca | -0.037 | H12 | ha | 0.149 |
| C25 | ca | -0.183 | H13 | ha | 0.160 |
| C26 | ca | -0.138 | H14 | ha | 0.160 |
| C27 | c3 | 0.020 | H15 | h4 | 0.218 |
| H1 | h1 | 0.054 | H16 | ha | 0.191 |
| H2 | hc | 0.074 | H17 | ha | 0.146 |
| H3 | hc | 0.074 | H18 | ha | 0.152 |
| H4 | hc | 0.037 | H19 | hn | 0.412 |
| H5 | hc | 0.037 | H20 | hn | 0.412 |
| H6 | h2 | 0.093 | H21 | hn | 0.300 |
| H7 | hc | 0.160 | | | |
| H8 | hc | 0.160 | | | |
| H9 | hx | 0.038 | | | |
| H10 | h1 | 0.084 | | | |
| H11 | h1 | 0.029 | | | |
| H12 | ho | 0.427 | | | |
| H13 | hc | 0.154 | | | |
| H14 | hc | 0.154 | | | |
| H15 | hc | 0.154 | | | |
| H16 | hn | 0.366 | | | |
| H17 | hn | 0.366 | | | |

| | | |
|---|---|---|
| H18 | h1 | -0.003 |
| H19 | h1 | -0.003 |
| H20 | ho | 0.452 |
| H21 | ho | 0.307 |
| H22 | ho | 0.396 |
| H23 | ha | 0.128 |
| H24 | ho | 0.410 |
| H25 | ha | 0.171 |
| H26 | ha | 0.146 |
| H27 | h1 | 0.061 |
| H28 | h1 | 0.061 |
| H29 | h1 | 0.061 |
| H30 | hn | 0.366 |

**Table S2:** Charge info of doxorubicin and niraparib

| Atom Type | σ (nm) | ε (kJ/mol) |
|---|---|---|
| ca | 0.3400 | 0.3598 |
| ha | 0.2600 | 0.0628 |
| ce | 0.3400 | 0.3598 |
| cf | 0.3400 | 0.3598 |
| c | 0.3400 | 0.3598 |
| o | 0.2960 | 0.8786 |
| oh | 0.3066 | 0.8803 |
| ho | 0.0000 | 0.0000 |
| os | 0.3000 | 0.7113 |
| c3 | 0.3400 | 0.4577 |
| h1 | 0.2471 | 0.0657 |
| n4 | 0.3250 | 0.7113 |
| hc | 0.2650 | 0.0657 |
| h2 | 0.2293 | 0.0657 |
| hx | 0.1960 | 0.0657 |
| hn | 0.1069 | 0.0657 |
| na | 0.3250 | 0.7113 |
| nc | 0.3250 | 0.7113 |
| n | 0.3250 | 0.7113 |
| cc | 0.3400 | 0.3598 |
| h4 | 0.2511 | 0.0628 |

**Table S3**: Atom types info for curcumin (an, enol and trans), doxorubicin and niraparib

| PURE CURCUMIN (kJ/mol) | | | | | |
|---|---|---|---|---|---|
| | **van der Waal** | **Electrostatic** | **Polar** | **SASA** | **Binding** |
| **Au+An** | | | | | |
| **IAU-CUR** | -3495.741 ± 32.203 | -1.903 ± 0.010 | -756.563 ± 34.318 | -24.191 ± 2.849 | -4278.397 ± 50.076 |
| **IAU-ETH** | -1411.330 ± 36.197 | 0.030 ± 0.013 | 71.114 ± 6.802 | 487.900 ± 17.061 | -852.286 ± 41.352 |
| **CUR-ETH** | -4458.23 ± 110.337 | -2397.093 ± 367.341 | 2567.147 ± 235.217 | -761.792 ± 15.279 | -5049.967 ± 358.975 |
| **Etotal** | -9365.3 | -2398.97 | 1881.7 | -298.08 | -10180.65 |
| **Au+Enol** | | | | | |
| **IAU-CUR** | -3689.008 ± 26.902 | -0.016 ± 0.005 | 138.655 ± 3.983 | -23.954 ± 3.048 | -3574.322 ± 26.847 |
| **IAU-ETH** | -1383.511 ± 26.179 | -0.001 ± 0.022 | 73.990 ± 9.385 | 493.946 ± 15.928 | -815.576 ± 29.748 |
| **CUR-ETH** | -3274.57 ± 126.278 | -806.638 ± 64.458 | 2818.741 ± 133.112 | -588.725 ± 16.731 | -1851.193 ± 83.885 |
| **Etotal** | -8347.09 | -806.66 | 3031.39 | -118.73 | -6241.09 |
| **Au+Trans** | | | | | |
| **IAU-CUR** | -4196.894 ± 30.410 | -0.015 ± 0.007 | 166.153 ± 5.546 | -32.324 ± 3.496 | -4063.080 ± 31.046 |
| **IAU-ETH** | -942.065 ± 21.421 | 0.003 ± 0.011 | 49.373 ± 6.271 | 498.084 ± 15.250 | -394.604 ± 26.822 |
| **CUR-ETH** | -3369.91 ± 153.526 | -1059.097 ± 88.211 | 3151.614 ± 142.302 | -635.942 ± 17.941 | -1913.338 ± 111.356 |
| **Etotal** | -8508.87 | -1059.11 | 3367.14 | -170.18 | -6371.02 |
| **MIXTURE CURCUMIN (kJ/mol)** | | | | | |
| | **van der Waal** | **Electrostatic** | **Polar** | **SASA** | **Binding** |
| **Au+An-Enol** | | | | | |
| **IAU-CUR** | -3740.327 ± 31.603 | -0.947 ± 0.009 | -225.845 ± 25.117 | -24.482 ± 3.422 | -3991.601 ± 39.197 |
| **IAU-ETH** | -1135.228 ± 22.091 | 0.026 ± 0.013 | 49.606 ± 4.688 | 495.107 ± 14.970 | -590.488 ± 27.744 |
| **CUR-ETH** | -3431.407 ± 77.832 | -1071.325 ± 194.601 | 2774.105 ± 158.447 | -630.006 ± 11.884 | -2358.634 ± 167.834 |
| **Etotal** | -8306.96 | -1072.25 | 2597.87 | -159.38 | -6940.72 |
| **Au+Trans-An** | | | | | |
| **IAU-CUR** | -4063.584 ± 34.981 | -1.032 ± 0.010 | -186.947 ± 18.950 | -25.933 ± 3.753 | -4277.496 ± 43.096 |
| **IAU-ETH** | -981.177 ± 23.676 | 0.030 ± 0.009 | 41.701 ± 6.167 | 505.709 ± 15.398 | -433.737 ± 30.906 |
| **CUR-ETH** | -3486.697 ± 79.555 | -1172.093 ± 207.130 | 2811.039 ± 152.790 | -636.757 ± 12.908 | -2484.507 ± 192.566 |
| **Etotal** | -8531.46 | -1173.1 | 2665.79 | -156.98 | -7195.74 |
| **Au+Trans-Enol** | | | | | |
| **IAU-CUR** | -3998.938 ± 32.391 | -0.005 ± 0.009 | 141.909 ± 4.817 | -25.236 ± 3.387 | -3882.270 ± 33.037 |
| **IAU-ETH** | -1027.733 ± 32.680 | 0.010 ± 0.009 | 59.571 ± 6.159 | 506.075 ± 13.359 | -462.076 ± 35.339 |
| **CUR-ETH** | -3399.082 ± 78.211 | -918.072 ± 62.612 | 3074.308 ± 82.643 | -625.883 ± 12.399 | -1868.728 ± 77.122 |
| **Etotal** | -8425.75 | -918.07 | 3275.79 | -145.04 | -6213.07 |

**Table S4:** MMPBSA values for Au+curcumin (pure and mixtures).

| MMPBSA (kJ/mol) – With ethanol | | | | | |
|---|---|---|---|---|---|
| **System** | **van der Waal** | **Electrostatic** | **Polar** | **SASA** | **Binding** |
| **Au+DOX** | -3264.74 ± 43.02 | 1.11 ± 0.02 | 78.83 ± 10.23 | -26.98 ± 3.45 | -3211.77 ± 44.22 |
| **Au+DOX+An** | -3298.16 ± 32.54 | -0.22 ± 0.01 | 131.12 ± 9.17 | -17.51 ± 3.71 | -3184.76 ± 34.28 |
| **Au+DOX+Enol** | -3047.16 ± 31.96 | 0.62 ± 0.01 | 98.74 ± 6.70 | -16.27 ± 3.41 | -2964.07 ± 32.58 |
| **Au+DOX+Trans** | -3483.50 ± 34.81 | 0.63 ± 0.02 | 132.36 ± 5.71 | -22.88 ± 3.63 | -3373.39 ± 35.62 |
| **MMPBSA (kJ/mol) – Without ethanol** | | | | | |
| **Au+DOX** | -3618.775 ± 57.021 | 1.168 ± 0.042 | 90.550 ± 14.920 | -30.101 ± 3.700 | -3557.158 ± 54.657 |
| **Au+DOX+An** | -3995.942 ± 32.056 | -0.244 ± 0.016 | 186.570 ± 17.159 | -32.298 ± 3.125 | -3841.914 ± 34.190 |
| **Au+DOX+Enol** | -4742.093 ± 32.407 | 0.000 ± 0.000 | 170.107 ± 13.840 | -28.633 ± 3.304 | -4600.620 ± 35.393 |
| **Au+DOX+Trans** | -4655.947 ± 51.459 | 0.709 ± 0.026 | 169.110 ± 11.249 | -31.260 ± 3.165 | -4517.388 ± 56.234 |
| **MMPBSA (kJ/mol) – Cancer membrane & Au+DOX+curcumin** | | | | | |
| **Au+DOX** | -678.159 ± 33.945 | -30513.455 ± 212.808 | 2892.350 ± 352.084 | -78.256 ± 3.391 | -28377.520 ± 413.850 |
| **Au+DOX+An** | -636.073 ± 79.691 | -844.366 ± 135.294 | 2073.896 ± 195.732 | -119.365 ± 7.267 | 474.093 ± 131.491 |
| **Au+DOX+Enol** | -433.892 ± 48.438 | -40087.256 ± 511.918 | 3557.146 ± 308.232 | -110.842 ± 7.443 | -37074.844 ± 304.260 |
| **Au+DOX+Trans** | -293.526 ± 26.482 | -38337.286 ± 282.891 | 1983.362 ± 283.224 | -79.184 ± 5.655 | -36726.634 ± 319.783 |

**Table S5:** MMPBSA values for between & curcumin+DOX.

| System | L-J Energy (kJ/mol) |
|---|---|
| Au+DOX | -453.336 |
| Au+An | -237.766 |
| Au+Enol | -253.339 |
| Au+Trans | -373.515 |

**Table S6:** L-J energy values between curcumin/DOX and gold nanoparticle.
